# A scalable method for modulating plant gene expression using a multispecies genomic model and protoplast-based massively parallel reporter assay

**DOI:** 10.1101/2024.12.05.626999

**Authors:** Sania Jevtic, Colleen Drapek, Christophe Gaillochet, Andrew Brockman, James Cadman, Timo Flesch, Nicolas Kral

## Abstract

Precision breeding tools such as CRISPR-Cas genome editing can speed up innovation in plant biotechnology and boost crop yields. The challenge remains to efficiently apply precision breeding methods to plant gene regulation. Endogenous gene regulatory sequences are subject to complex transcriptional control, a bottleneck in altering gene expression patterns in a predictable way. Here we present the CRE.AI.TIVE platform, enabling upregulation of plant gene activity without *a priori* knowledge of individual cis-regulatory elements or their specific location. A predictive machine learning model underpinning the platform has been trained on a wide range of tissue specific transcriptomic and epigenomic coverage datasets from DNA sequence of 12 plant species, showing competitive performance on RNA-seq coverage prediction across all species. Our platform further combines *in silico* DNA sequence mutagenesis and a protoplast-based massively parallel reporter assay (MPRA). We demonstrate the platform’s functionality by mutagenesis of a proximal promoter of the tomato gene SlbHLH96 which yields predictions of variant gene activity *in silico*. 2,000 sequence candidates with varying predicted gene expression strength were validated with MPRA in plant protoplasts, identifying variants with significantly upregulated gene activity. A portion of functional sequence variants were further individually evaluated with a fluorescence reporter assay and were observed to contain a new order of known cis-regulatory elements. The CRE.AI.TIVE platform offers a first-of-its-kind scalable method of gene upregulation in plants with native DNA sequences without the need for CRE cataloguing and rational promoter design.

## Introduction

A major increase in crop yields is needed to sustain a growing world population and provide food security. A promising way to achieve this is through innovative genetic approaches that improve crop resilience with more predictable outcomes [1]. Adoption of CRISPR-Cas genome editing technology for precision breeding in crops is an effective way to improve crop traits and provides an easier path to commercialisation in several jurisdictions over conventional genetic modification [2].

Of particular interest is the use of CRISPR-Cas technology to modulate gene expression of native plant genes. This approach enables more subtle phenotypic changes by affecting specific tissues, developmental stages or environmental responses without the negative pleiotropic effects associated with protein coding sequence changes [3, 4] or with the use of cisgenic or transgenic constructs [5]. While CRISPR-Cas modifications in gene regulatory sequences offer more fine-tuned phenotypes [6, 7] the consequences of such changes are more difficult to discern than if protein coding sequences are modified. Gene regulatory sequence changes can modify myriad complexities of transcriptional control, including tissue specificity, transcriptional redundancy and modularity of cis-regulatory elements (CREs). Other changes modify the spacing of CREs, chromosomal interactions, epistasis and CRE compensation, all of which modify gene regulatory output [8]. Interpreting this complexity limits the deployment of genome editing technology for modulating expression of individual genes. As a result, most examples of promoter genome editing result in a controlled downregulation of specific genes [6, 9].

The few examples of CRISPR-Cas applications for gene expression upregulation have relied on random promoter editing [6, 10–12] and mutagenic events with large potential impact on the overall cell metabolism such as the removal of a gene between an active promoter and the protein coding sequence [13] or, even larger, chromosome-scale modifications [14]. While the application of CRISPR knock-in technology in plants continuously improves [15–17], the rational design of promoters is dependent on crop-specific experimental identification of individual CREs with non-generalisable logic [18]. Furthermore it relies on experimental assessment of CRE positions within a gene regulatory region and further experimental validation of gene regulatory activity for individual target applications, which presents an obstacle to scalability [18, 19].

Parallel to the progress of plant biotechnology and the adoption of genome editing has been equally rapid adoption of deep learning methods applied to genomic data [20–38]. The complexity associated with transcriptional control and identification of gene regulatory sequence determinants together with the influx of data availability fits well with end-to-end deep learning approaches. A number of models based on human data have addressed the challenge of predicting quantitative gene expression, measured using RNA-seq, directly from DNA sequence [25, 35, 39]. Borzoi [25], a (UNet) convolutional neural network (CNN) - transformer model, used 524 kilobase pair (kbp) sequences to predict thousands of epigenomic and transcriptomic coverage tracks in human and mouse. It exhibited impressive performance on the gene expression task and was able to recover the impact of known gene variants on gene expression. Further analysis of the model, including an independent study [40], revealed the ability to detect CREs and cell type specific enhancers.

In plants, CNNs have been employed for gene expression classification from relatively short sequences (promoter and terminator segments up to 3 kbp) by using either one, two or four plant species genomes [20–22]. Scaling up, the authors of [24] used masked language modelling to train AgroNT, a BERT-style transformer [41], on 48 plant species using 6 kbp sequences. The model was evaluated on a quantitative gene expression prediction task and displayed moderate performance. Hence, the complexity of plant genomics still poses a challenge for efficient design of new crop traits using existing machine learning models.

To overcome the limitations associated with rational promoter design and generate CRISPR-Cas applicable upregulating sequences, we present the CRE.AI.TIVE platform. This method combines a machine learning trained to predict transcriptomic and epigenomic coverage tracks from plant genome sequence data, iterative *in silico* gene regulatory sequence mutagenesis and protoplast-based gene expression validation. To demonstrate its functionality, we evolved novel tomato sequence variants for the proximal promoter of the gene SlbHLH96, previously observed to contribute towards tomato drought tolerance with transgenic upregulation [42]. Gene regulatory activity of 2,000 novel sequences was measured in a protoplast-based massively parallel reporter assay (MPRA), an approach outlined in [43, 44]. Additional validation of select variants demonstrates increased gene expression with fluorescence.

## Results

### The CRE.AI.TIVE model accurately predicts plant gene expression

The CRE.AI.TIVE model is a machine learning model, inspired by the Enformer and Borzoi models [24, 25], with a CNN-transformer architecture that is trained to simulate cell transcription (Fig. 1A). Our model accepts a plant DNA sequence with length 2^16^ = 65, 536 bp as its input and its output target is a prediction for a transcriptomic or epigenomic coverage track every 32 bp along the sequence from a species-specific output head. Model training is done in two stages: pre-training and fine-tuning. The two-step training strategy makes our solution flexible and scalable since pre-training is resource-heavy but only needs to be done once. On the other hand, fine-tuning can be achieved relatively rapidly and, by virtue of transfer learning, results in a performant model even when the amount of fine-tuning data is scarce.

**Figure 1:**
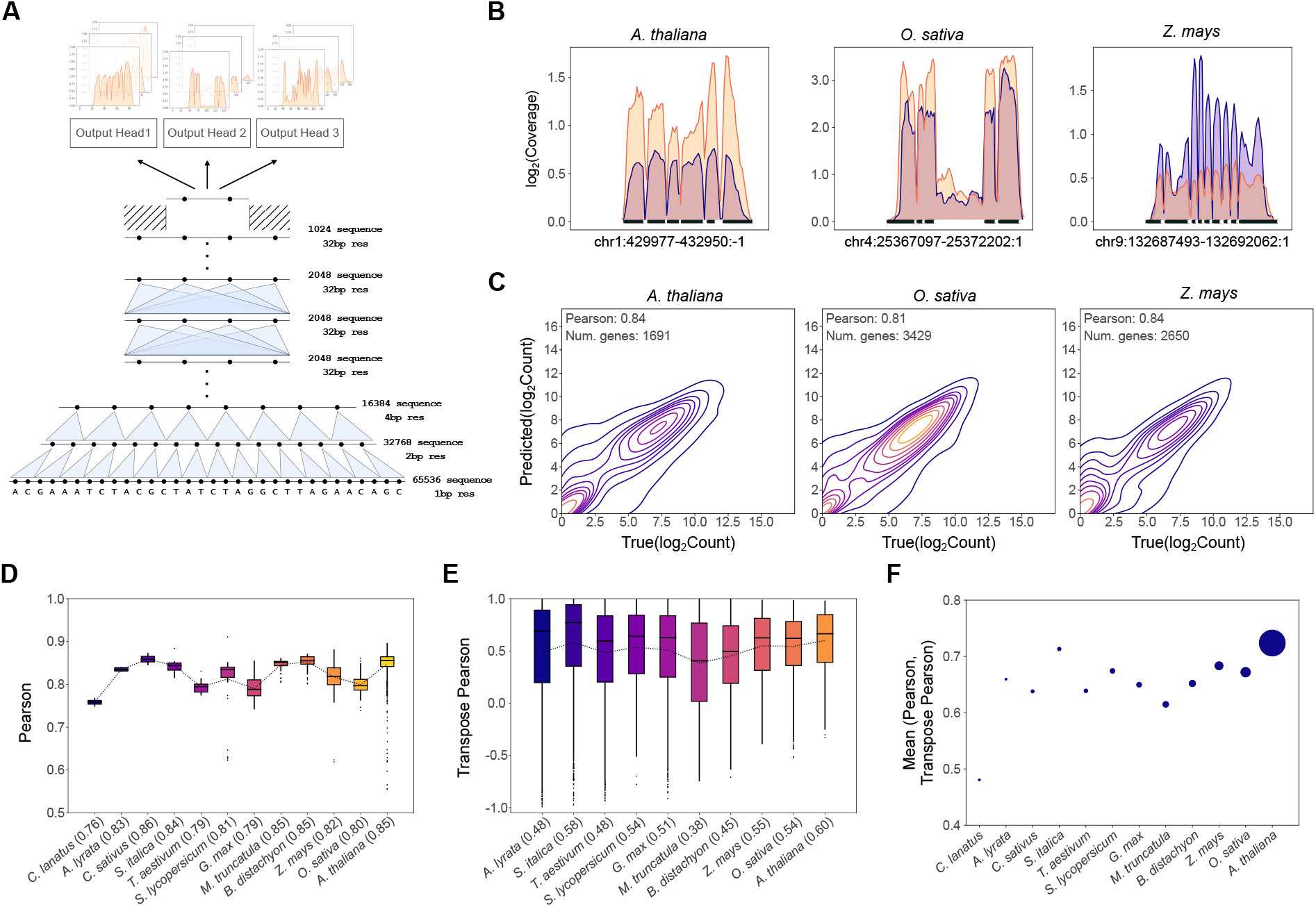
The CRE.AI.TIVE model and its performance on unseen data. **(A)** The model architecture: convolutional layers downsample the 65 kbp input sequence (bottom) before passing through transformer layers. Species-specific output heads (top) predict coverage tracks at 32 bp resolution. **(B)** True (purple) and predicted (orange) RNA-seq coverage for a selection of species, genes and experimental conditions: (left) *Arabidopsis thaliana* gene AT1G02220, leaf grown in short-day condition [45], (middle) *Oryza sativa* gene Os04g0507700, rice seedling [46], and (right) *Zea mays* gene Zm00001eb394070, shoot apical meristem 19 days after germination grown in a greenhouse large pot [47]. Genes can come from either strand but are forward oriented in the plots. Exons are displayed in thick black on the x-axis. **(C)** Contour plots of predicted vs true gene expression log_2_(count) values for the same species and experimental conditions as Fig. 1B, but with all validation genes shown. The Pearson correlation coefficient is displayed. **(D)** Box plot distributions of Pearson correlation coefficients between predicted and true log_2_(count) values conditional on RNA-seq targets. The dotted line joins the box means, which are also indicated in brackets next to species names along the bottom. Species are ordered according to the total number of targets (transcriptomic plus epigenomic). **(E)** Same data shown in (D) but as Pearson correlation coefficients conditional on genes (aka the “transpose Pearsons”, see Methods). *Cucumis sativus* and *Citrullus lanatus* are omitted since transpose Pearsons are calculated across targets and these species only have two targets. **(F)** The average of Pearson and transpose Pearson values for each species. The sizes of the dots correspond to the total number of targets.

Pre-training is performed on a large dataset composed of multiple species to learn regulatory signals that are shared across plants. In the present case, we pre-trained the model on 12 plant species, shown in Table S1 (Supplementary Material). When comparing the experimental (“ground truth”) vs model-predicted RNA-seq coverage for a selection of genes, the model recovers the shape of the RNA-seq coverage signal, especially at the boundaries of the peaks and troughs (Fig. 1B). This suggests that the model has learned the sequence motifs that signify intron-exon boundaries. Additional coverage plots are provided in Fig. S1. To measure the ability to predict gene expression from RNA-seq coverage we computed a gene-level count for the 12 species (Fig. 1C, Fig. S2). A general pattern is observed, namely, that the model is able to capture higher expression values well, but struggles to predict silent genes to some extent, when true expression is zero. In order to assess the model predictive ability, we calculated the distributions of Pearson values of log_2_ gene expression counts conditional on RNA-seq conditions, separately for each species (Fig. 1D). The model displays competitive performance: the all-species average Pearson is 0.82 and the values are distributed relatively narrowly despite the fact that: (i) with the exception of the model plant *Arabidopsis thaliana*, all plant species have far fewer target datasets than Borzoi; (ii) the number of points (the RNA-seq target conditions) in each box varies very widely, (iii) the number of validation genes varies between species, and (iv) the dynamic range of the gene expression count values is large, as seen from the gene expression count plots Fig. 1C.

We then assessed the model by computing a “transpose Pearson” (Fig. 1E) which represents the Pearson correlation values conditional on the genes (see Methods). With an all-species average of 0.48, these are lower than Pearson’s correlation in Fig. 1D and have a wider distribution, implying that predicting variation between RNA-seq conditions is harder than between genes - a trend similar to that observed in Borzoi [25]. However, the model explains a substantial amount of the variation in both genes and RNA-seq conditions. Since both perspectives are important, for each species we computed a summary statistic as the average of the Pearson and transpose Pearson, and compared this against the total number of target conditions (transcriptomic and epigenomic) (Fig. 1F). The two quantities are correlated hence outlining a strategy for further improving performance. While the amount of target data for most species is an order of magnitude less than the model species *Arabidopsis thaliana*, the model still exhibits convincing predictive capabilities across all species, which is due to mixing species during pretraining. This strongly indicates a level of conservation of important biological features such as CREs across plant species, with model performance for underrepresented species being bolstered by signals learned from the model species.

### The CRE.AI.TIVE model can be fine-tuned with cultivar-specific data and used to guide directed mutagenesis of select genes

We fine-tuned the pre-trained model to predict gene expression for *Solanum lycopersicum* cv. Ailsa Craig, our experimental cultivar, using in house data. This cultivar is different to the one seen by the model during pre-training, which is cv. Heinz 1706. For fine-tuning, we introduced an output head for the prediction of gene expression in fragments per kilobase million (FPKM) units of four tissues: cotyledon, leaf, root, stem. The Pearson and transpose Pearson values for cv. Ailsa Craig FPKM predictions are 0.80 and 0.46 on unseen data across the genes and tissues respectively, similar to the *Solanum lycopersicum* cv. Heinz 1706 pre-trained prediction performance: 0.81 and 0.54 (Figs. 1D and E), indicating that the model can accurately predict gene expression for cv. Ailsa Craig.

Our *in silico* mutagenesis algorithm predicts sequence perturbations that achieve a particular level of predicted gene expression (see Methods). As such, the model is used as a predictor of gene expression in the mutagenesis process. We aimed to increase the expression of SlbHLH96 (gene ID Solyc11g056650.2) in the cotyledon tissue. We first assessed the model’s predictive ability on SlbHLH96 across four tissues (Fig. 2A and Fig. 2B). The predictions were conducted by the all-splits trained model, hence the model has seen the wild-type DNA sequence during training (see Methods). We then subjected the wild-type DNA sequence to 100 mutagenesis iterations, with each generation replacing, adding or removing a single base pair only in a 211 bp region of the proximal promoter (Fig. 3A). Millions of variants were tested during this process, but only 2,000 sequence variants representing a range of generations with varying predicted gene expression strengths (Fig. 2C) were subsequently selected for experimental validation. We observed predicted transcriptional activity to amplify across the whole coding sequence, increasing with further sequence perturbations (Fig. 2D).

**Figure 2:**
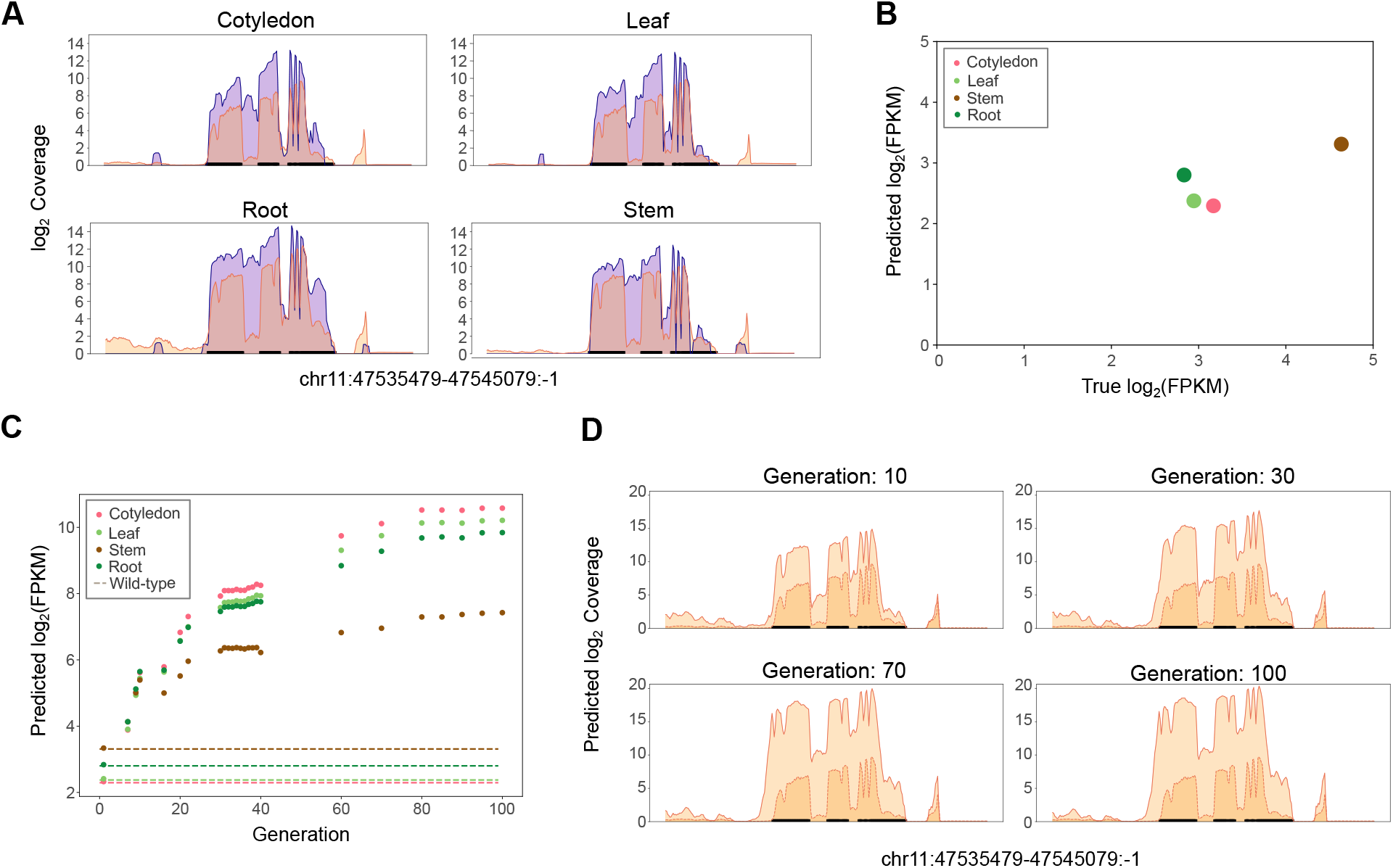
Solanum lycopersicum cv. Ailsa Craig SlbHLH96 (Solyc11g056650.2) prediction and mutagenesis. **(A)** Coverage of the cv. Ailsa Craig SlbHLH96 gene for the four fine-tuned RNA-seq tissues, showing the ground truth (purple) and predicted (orange) values. **(B)** Predicted vs true SlbHLH96 gene expression values (in log_2_ FPKM units). **(C)** Evolving the wild-type SlbHLH96 sequence using our mutagenesis algorithm increases mean generation predicted expression over the wild type (dashed line). **(D)** Predicted cotyledon coverage of mutants (solid line) from generations 10, 30, 70 and 100 versus wild type (dashed line).

**Figure 3:**
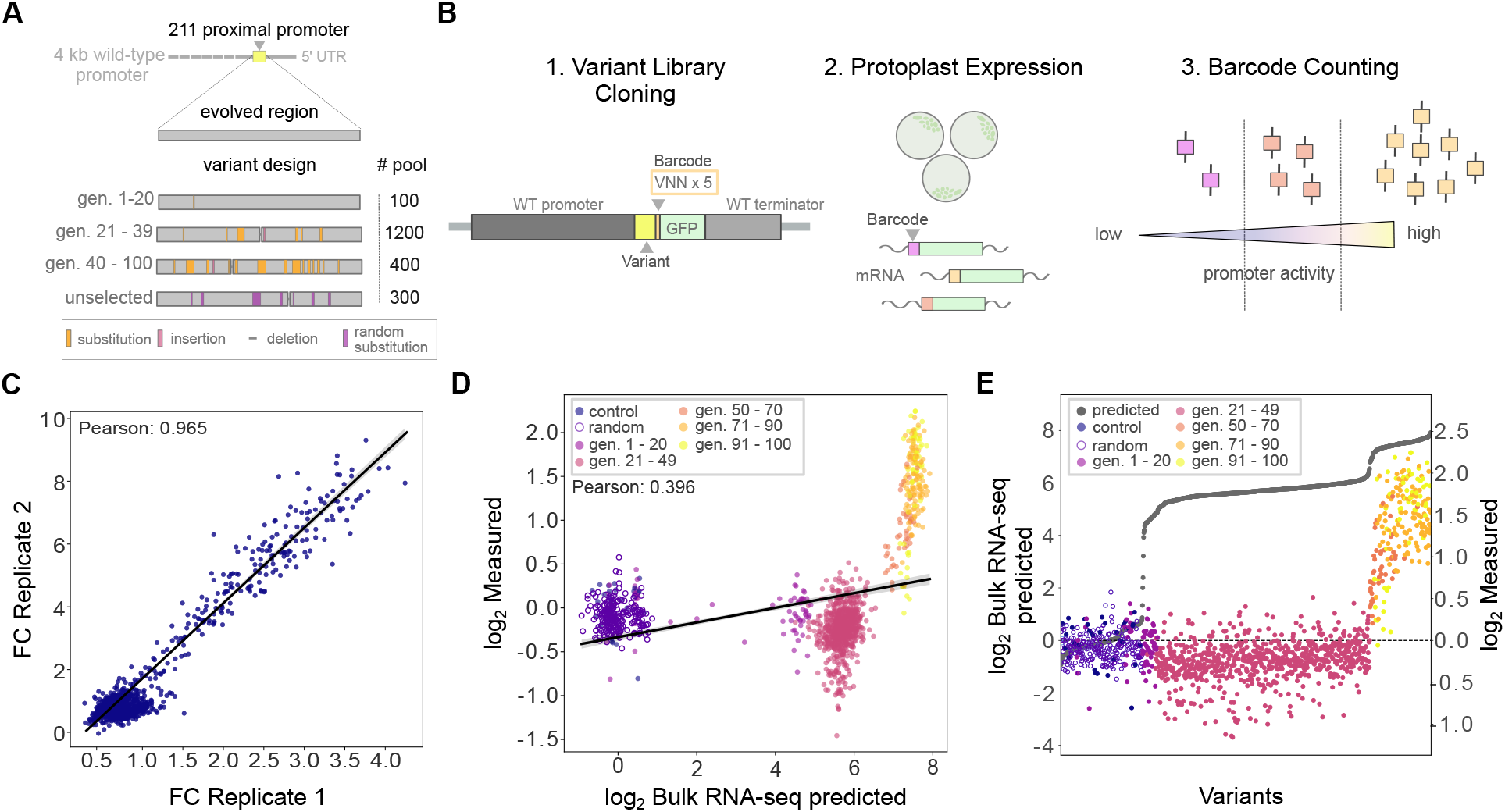
An MPRA to test CRE.AI.TIVE in tomato protoplasts. **(A)** A schematic of the selection of evolved proximal promoters for testing in the MPRA. Variants were selected from a range of generations and represented in different amounts. Examples of types of DNA changes (substitutions, insertions, and deletions) are shown for illustrative purposes. Unselected sequences from generations 13-50 are evolved by *in silico* mutagenesis but not selected for gene expression (i.e. random sequences). **(B)** A schematic of the MPRA library backbone construct. Each variant resides downstream a wild-type promoter fragment and upstream a 35S 5’UTR, a start codon, a 15 bp barcode (VNNx5) and a GFP CDS. A wild-type 3’ UTR and terminator sequence flanks the 3’ end of the GFP coding sequence. The library is transfected into protoplasts and RNA is harvested for barcode counting and normalization. **(C)** A plot comparing the normalized RNA/DNA ratio of replicate 1 (x-axis) and replicate 2 (y-axis). The Pearson correlation is 0.965. Black line shows the line of best fit. **(D)** A plot comparing the average measured fold change across replicates (x-axis) to the CRE.AI.TIVE bulk RNAseq predicted value. Black line shows the line of best fit. The Pearson correlation is 0.396 **(E)** A plot comparing the measured and predicted fold change values (y-axis) by variant (x-axis). Point color indicates generation of evolution, random sequence or control.

### A massively parallel reporter assay in tomato protoplasts reveals CRE.AI.TIVE designs a range of promoter activities

In order to measure evolved promoter sequence activity, we selected 2,000 proximal promoter sequences with a range of predicted activities to test in a massively parallel reporter assay (MPRA) in tomato protoplasts (Fig. 3A, see methods). Specifically, we selected 100 from generations 1-20, 1,200 from generations 21-39, 400 from generations 40-100 and 300 random sequences (13-50 undirected mutagenesis generations). We constructed a library of proximal promoter fragments approximately 211 bp with the aim to evaluate their function in a relevant genomic context. The ∼211 bp evolved sequence was inserted into a plasmid downstream a cloned fragment of the native 3.6 kbp SlbHLH96 promoter and upstream a barcode sequence, a GFP coding sequence and a cloned fragment containing the 2 kbp 3’UTR and terminator region of the native SlbHLH96 genomic region (Fig. 3B).

The library was transfected into tomato cotyledon protoplasts and harvested after 24 hours for RNA isolation. Promoter activity was measured using a tailored version of MPRAflow [44] and MPRAnalyze [48] to normalize barcode counts and compute an RNA/DNA ratio (Fig. 3B). All variants not represented by at least 10 barcodes were removed from the analysis, leaving a total of 1,304 variants for downstream analysis. Between experimental replicates there was a Pearson correlation of 0.965 (Fig. 3C) indicating high reproducibility of measured promoter activity in our library.

We next asked how the measured fold change compares to the predicted activity (Fig 3D). We found the model predictions have a positive correlation with the measured activity, with a Pearson’s correlation of 0.396. The model more accurately predicts later generations of evolved variants than earlier generations, which could reflect either limitations of the model or in comparing MPRA and RNA-seq data (see Discussion, [36, 43]). In order to better understand how each individual variant performs, we compared the measured activity to the predicted activity for each variant (Fig. 3E). A pattern emerged that the CRE.AI.TIVE can predict the activity of the SlbHLH96 variants after several generations of evolution (Fig. 3E). It also captures transcriptional activity increase in a shape typical of cooperative activity, a common behavior in transcription factor binding [49, 50].

Of the 1,700 promoters that we screened for predicted increase in transcriptional activity, we validated 163 promoters with at least two fold increase in activity comparison to controls, 32 of which have a four fold increase in comparison to controls. This demonstrates that the CRE.AI.TIVE platform is a useful tool for modulating promoter strength.

### CRE.AI.TIVE designs functional promoters for overexpression by creating transcription factor motifs

Following our MPRA results, we selected 3 variants with the highest level of measured promoter activity for follow up analysis and further validation in protoplasts. These variants were cloned in a similar fashion to Fig. 3B but lacked the barcode sequence before the GFP coding sequence. In protoplasts, variants were visibly brighter compared to wild-type controls (Fig. 4A). We quantified the fluorescence of protoplasts and found all variants had distributions significantly brighter than the wild-type control (Fig. 4B). We further analysed cells by Fluorescence Activated Cell Sorting (FACS) comparing the brightness of GFP-positive cells in protoplasts transfected with variants or control constructs. We found that all variants have a 1.5 - 2 fold higher GFP brightness than wild-type, with higher frequencies of cells within the GFP detection range (Fig. 4C). In general, the selected promoter variants show increases in RNA/DNA ratio are related to increases in expression of GFP, supporting the use of CRE.AI.TIVE for changing protein levels by modulating promoter activity.

**Figure 4:**
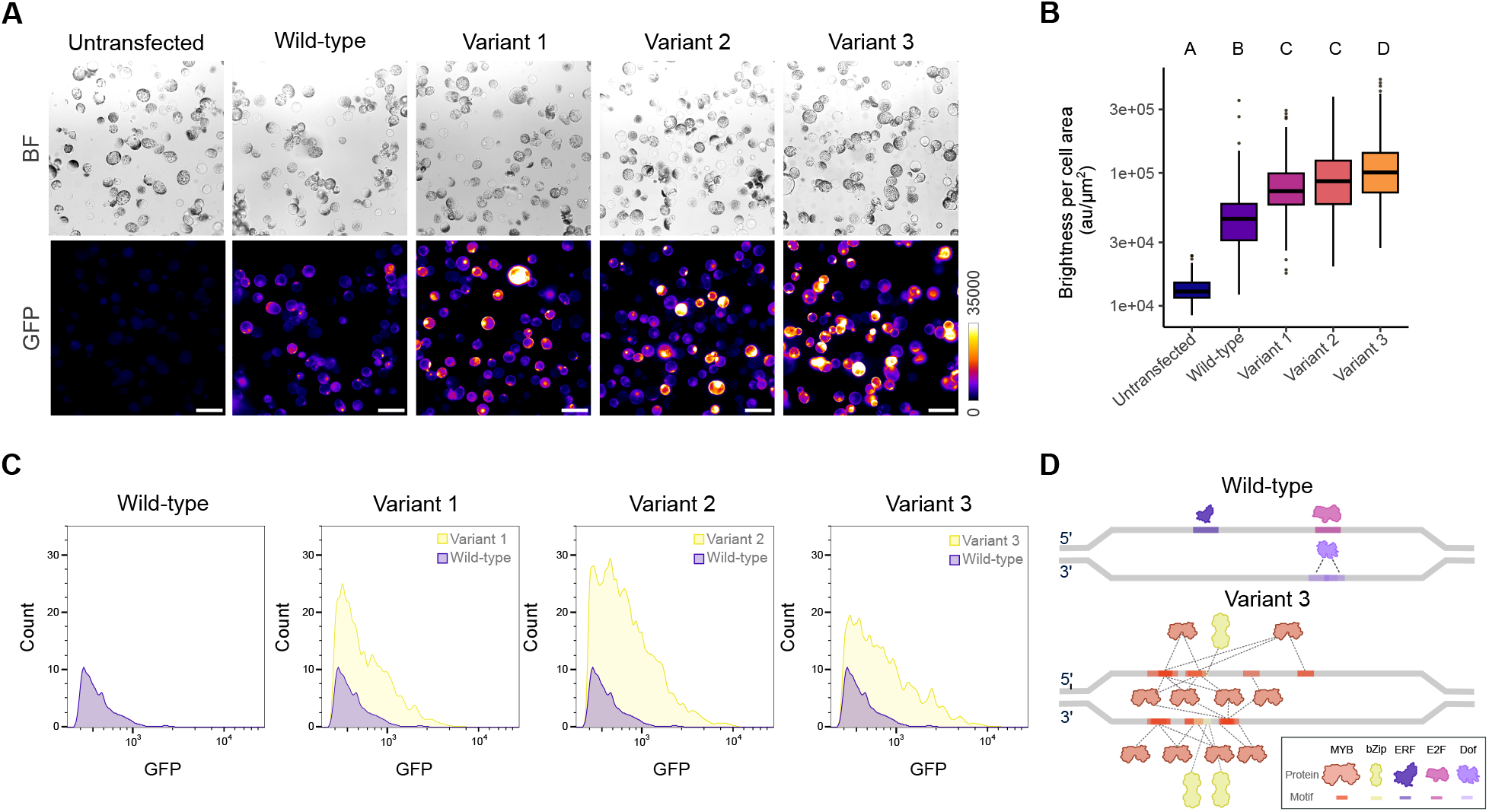
CRE.AI.TIVE generates functional overexpressing promoters using known transcription factor motifs. **(A)** Brightfield images and GFP fluorescence in protoplasts transfected with the wild-type promoter, top variants, and untransfected control. Scale bar =100 µm. **(B)** Quantification of fluorescence in protoplasts transfected with the wild-type promoter, top variants, and untransfected control. Different letters above the plot indicate significantly-different fluorescence distributions as determined by Kruskal-Wallis test followed by a Dunn’s test with Bonferroni adjustment. **(C)** Histograms of FACS-analyzed protoplasts transfected with the wild-type promoter and top variants. Results from the wild-type promoter are represented in each plot for comparison. **(D)** Schematic representing the FIMO-identified transcription factor motifs in the wild-type promoter and in variant 3. Each dashed line indicates a predicted binding interaction.

Given the demonstrated function of these promoters, we next interrogated variant sequence structure to identify sequence changes that could confer higher gene expression. We searched the wild-type sequence for identified tomato transcription factor binding sites using the Find Individual Motif Occurrences (FIMO) with *S. lycopersicum* binding motifs downloaded from the Plant Transcription Factor Database [51, 52]. In the wild-type proximal promoter, 4 transcription factor motifs were identified - 1 from the ethylene responsive factor (ERF) family, 1 from the E2F family and 2 from the Dof zinc finger family (Fig. 4D, Table S2). To determine if there were changes in binding motifs, we focused on promoter variants with a fold change greater than 4 (n = 32) and narrowed our analysis to variant 3, the promoter with the highest GFP expression in protoplasts. This variant gained 97 substitutions and 3 deletions, for a total of 100 bp changed in comparison to the wild-type sequence. Using a FIMO search on variant 3, we identified 26 overlapping motifs from 13 transcription factors, 11 of which are in the MYB family and 2 that are in the bZip family (Fig. 4D, Table S2). Variant 3 has gained a new order of known transcription factor binding motifs, driving the change in observed expression.

## Discussion

Machine learning approaches have enabled significant strides in understanding principles of biology. This is especially the case in protein design and engineering where models like AlphaFold [53] and its subsequent versions have enabled rapid protein structure research and bespoke design to suit a breadth of applications for discovery and human health. Here, we present an important step towards gene regulation design in plants to upregulate expression for a specific application in drought resilience. We present CRE.AI.TIVE, a platform based on principles that include deep learning-based gene expression prediction from DNA sequence and in protoplast MPRA screening. Using this approach we designed and validated thousands of SlbHLH96 proximal promoters with varying gene expression strength. A selection of these demonstrated upregulated promoter activity, with 32 variants having higher than four fold activity.

The utility of the CRE.AI.TIVE platform presents solutions to the challenges that are limiting progress of gene regulation engineering in plants. While we focused on one application, this approach is in principle scalable across all the genes in the 12 crop species present in the model training dataset and does not rely on manual, individual CRE validation, or any other form of genomic annotation. This approach also presents an achievement in terms of engineering higher gene activity from endogenous promoters, an important shift away from the use of viral and other synthetic promoters that can cause detrimental pleiotropic effects [54, 55]. Identifying the modifications that enable overexpression of native genes in plants is particularly relevant for precise genome editing with CRISPR-Cas technologies.

Traditionally, identifying the function of promoter elements has been slow, labour intensive and frequently ungeneralizable across species. Remarkably, we show that, in addition to designing a functional overexpressing promoter, the deep learning model has learned regulatory logic without a priori knowledge of known transcription factor motifs. The platform output is a proximal promoter with changed overlapping transcription factor binding motifs of several transcription factors, a strategy described in other plant ubiquitous promoters as optimizing expression for many possible tissue contexts [56]. We speculate that this approach towards promoter engineering may also aid with other aspects of promoter structure that affect gene regulation. In the future, we plan to evaluate the performance of CRE.AI.TIVE in learning rules relating to motif flanking sequences and motif positioning in reference to each other and to transcription start sites as CRE positioning is a major determinant of gene regulatory output [57–59].

Challenges facing further improvement of the CRE.AI.TIVE platform remain. Of these, a major challenge is data annotation and availability. In comparison to the large amount of RNA-seq and active chromatin data that is available for human and mouse, a smaller set of data exists for individual plant species with the majority in model species such as *Arabidopsis thaliana*. Additionally, while the model makes accurate predictions for bulk RNA-seq data, the correlation between protoplast MPRA data and model predictions for individual variants of a mutagenised promoter are lower, though the correlation does improve with increasing mutagenesis generations. Partially this is because the model was not trained on RNA-seq data collected from tomato protoplasts, and a discrepancy between bulk RNA-seq and MPRA is already present at the experimental level [36, 43]. However it was also observed that coverage models do have lower accuracy on perturbation data, especially involving perturbations in distal parts of promoters [60]. Further work also remains in validating the activity of evolved SlbHLH96 promoter variants in tomato plants and its contribution towards drought tolerance.

Beyond upregulation, a future application of the CRE.AI.TIVE platform is towards modulating gene sequences for more tissue-specific, temporal or stress-induced expression. While protoplasts offer scalability and flexibility for rapidly testing gene expression, validation of gene activity in plants will be a key step beyond modulating for constitutive overexpression. Ultimately, modifications in native genes in plants are necessarily dependent on strides in CRISPR-Cas technology improvements for precise genome editing to introduce these changes. Developments towards specific promoter editing technology would be invaluable for tackling challenges facing agriculture in a rapidly changing climate.

In summary, the CRE.AI.TIVE platform designs and validates plant promoters without a priori knowledge of individual motif sequences or their localisation and structure. We have demonstrated the utility of this platform in designing a range of SlbHLH96 promoters with different gene expression strengths that could be used for enhancing tomato drought tolerance.

## Methods

### The CRE.AI.TIVE model pre-training dataset

To model transcription, the target transcriptomic data consisted of RNA-seq coverage tracks. In addition, epigenomic data including ChIP-seq, ATAC-seq, DNAse-seq, FAIRE-seq and DAP-seq experiments were included, where available, as they encode gene regulatory information and aid in understanding tissue specificity.

The pre-training dataset leveraged data from public sources. The genomic sequence and annotation data came from reference genomes downloaded from the Ensembl Plants release-56 [61]. Sequences were centred on transcription start sites (TSS) of the plant’s genes, and have a context window of 32,768 bp either side of the central TSS, resulting in a total sequence length of 65,536 bp.

Epigenomic data was obtained from ChIP-Hub [62], which hosts a large database of pre-aligned raw reads and curated metadata for ChIP-seq, ATAC-seq, DNAse-seq, FAIRE-seq or DAP-seq experiments.

To help select the appropriate transcriptomic RNA-seq experiments, we used curated metadata from PlantExp [63]. To ensure the model learns the regulatory patterns of native genes, we chose RNA-seq experiments performed on the “wild-type” genotype only and a cultivar matching that of the reference genome. Applying these constraints to the metadata resulted in 12 target plant species.

Table S1 (Supplemental Material) presents the number of samples we sourced for each species and experiment type. No epigenomic samples were available for *Citrullus lanatus, Solanum lycopersicum*, and *Zea mays* since the reference genome versions used by ChIP-Hub for alignment of these three species did not match Ensembl Plants release [61]. Biological replicate samples for each experimental condition are aggregated into target conditions to marginalize out biological replicate variation before model training. The number of these resulting targets is also shown in the table.

We generated RNA-seq coverage data from raw RNA-seq reads by alignment to corresponding reference genomes. Raw reads were downloaded from the open access NCBI SRA AWS S3 bucket [64]. Parallelised RNA-seq alignment was done via the AWS HealthOmics running a customised version of the nfcore/rnaseq pipeline [65] with HISAT2 [66] as the alignment algorithm and deepTools [67] bamCoverage to produce the coverage BigWig files. The sample coverage data was normalised to RPKM count units and replicates were aggregated.

The training data is separated into train, validation and test splits, allocating approximately 80%, 10%, and 10% to the train, validation and test splits respectively. The split itself is implemented in a way that encourages homologous genes to reside in the same split and input sequence. This is to reduce possible information leakage between splits due to evolutionarily related genes [21, 24, 25, 38], which can occur either within or between species. The Ensembl Plants release-56 gene trees were used to define the homology and a split was deduced for its species.

### The CRE.AI.TIVE model

Length 2^16^ bp = 65, 536 bp input DNA sequences are first one-hot encoded into a 4-dimensional representation before passing through five convolutional neural network layers that build up a local representation of sequence motifs in their filters. Each convolution applies a pooling of size 2 which results in downsampling to a resolution of 32 bp at each spatial coordinate. This choice was a compromise between retaining granular enough DNA sequence information to resolve exons versus the growing time and memory dependence of the subsequent transformer layer on sequence length. Following convolution, long distance interactions between sequence features are achieved by a stack of transformer units using multi-headed attention. After this, only the middle half of the sequence is retained by cropping 25% from each binned sequence edge. This is to mitigate against poor predictions that can occur if the amount of signal available from either side of a spatial target prediction is imbalanced. The cropped transformer outputs are fed through a series of output heads, resulting in a prediction of 32 bp binned coverage signal for each target condition. A separate output head is allocated to each species and each transcriptomic or epigenomic category, generating values for multiple experimental conditions simultaneously at each 32 bp binned spatial coordinate. The resulting model has 140 million parameters. The model was trained using a mean squared error objective on log_2_ transformed target counts, with each output head contributing to the total loss. The model was trained using an AWS EC2 p5 instance powered by 8 H100 NVIDIA GPUs, with pre-training taking 2 weeks. The model and training code was implemented in PyTorch.

### Performance measures

The same method as Borzoi [25] was employed to obtain gene-level expression counts by summing RNA-seq target coverage in bins overlapping exon annotations. This unit is referred to as the gene expression count. A pseudo-count of +1 is added before applying a log_2_ transformation to the count; this is also applied to FPKM units.

From the gene expression log_2_(count) values, two metrics can be computed in order to obtain a summary of model performance per plant species: 1. a Pearson’s correlation value for accuracy of prediction across genes conditional on a target, and 2. a Pearson’s correlation value for accuracy of prediction across targets conditional on a gene. The second metric is referred to as the “transpose Pearson”.

### Mutagenesis

Mutagenesis is a probabilistic search algorithm that iteratively evolves sequences, beginning with a wild-type gene and successively accumulating mutations over a series of generations. In each generation, an individual insertion, deletion or substitution is introduced and the model predicts a target value for each mutated sequence. After inference, a subset of mutated sequences is sampled and these become the inputs for the next generation of mutagenesis. The procedure was set up in a tissue-specific way for mutagenesis of SlbHLH96 in order to maximise expression in leaf tissues, while keeping other tissue expression values constant. Since mutagenised sequences can be classified as unseen by the model, to achieve highest accuracy predictions, the model is trained on all splits before mutagenesis.

### Whole Genome Sequencing of Cultivar Ailsa Craig

Cultivar Ailsa Craig was grown to 7 weeks and DNA was extracted from leaf following standard DNA extraction protocols. The DNA was processed with a KAPA Hyper Prep PCR Free Kit (Roche), and sequenced with Illumina NovaSeq 6000 with PE150 reads, generating 413,437,952 reads. The reads were aligned to the SL3.0 reference genome using BWA [68] and SAM tools [69].

### Transcriptome Sequencing of Cultivar Ailsa Craig

RNA was extracted from 7 day old cotyledons, and 7 week old root, stem and leaf tissues, with 4-5 biological replicates each, according to standard RNA extraction protocols. The RNA was processed with the IDT xGen Broad-Range RNA Library kit, and sequenced with Illumina NovaSeq 6000 with PE 150 reads, generating a total of 841,750,716 reads across all samples. Similarly to publicly sourced data, reads were aligned to SL3.0 reference genome via the AWS HealthOmics running a customised version of the nfcore/rnaseq pipeline [65] with HISAT2 [66] as the alignment algorithm and deepTools [67] bamCoverage to produce the coverage BigWig files. The sample coverage data was normalised to RPKM count units and replicates were aggregated. The StringTie programme [70–73] was used to compute gene-level FPKM values.

### Massively Parallel Reporter Assay

#### MPRA library cloning

The SlbHLH96 (Solyc11g056650.2) genomic sequence was acquired from tomato cv. Ailsa Craig. A region of 211 bp upstream of the transcription start site was selected for evolution. The size of the variant region is constrained by the maximum length of amplicons for subsequent barcode-variant association NGS. We cloned a genomic fragment 3,608 bp upstream of the 211 proximal promoter region. Sanger and full plasmid sequencing revealed 4 substitutions and 2 deletions in the 3,608 bp region differing from the reference genome. The genomic region downstream the SlbHLH96 stop codon was not mapped when using the above method, so the sequence was acquired from TS9 genome from the Tomato Graph Pangenome project, sourced from Sol Genomics Network [74, 75]. We cloned a 2,002 bp downstream region including the 3’UTR and terminator. Fragments were subcloned into a vector backbone containing the promoter, variant entry site, GFP and terminator using the GreenGate system [76].

An oligo pool of 2,000 variants was provided by Twist Biosciences. Variants contained ∼211 bp evolved sequence flanked by cloning adapters. The 15 bp degenerate sequence (VNNx5) and 35S 5’UTR are added in pool amplification. Barcoded variant molecules were introduced into the library backbone using NEB HiFi Assembly. The library complexity was bottlenecked at ∼165 barcodes per variant.

#### MPRA library expression

The library was transfected into tomato protoplasts (10 *µ*g per 100K cells) following the protocol established by Nicolia et al [77]. Total RNA was harvested from two separate transfection experiments. RNA and DNA counts were generated as described in [44]. Amplicons for RNA counts and DNA counts were sent for Illumina 150 bp PE sequencing (Novogene).

The barcode association library was generated by amplifying the plasmid library with PCR primers upstream the variant region and downstream the barcode. Amplicons were sent for Illumina 250 bp PE sequencing (Novogene). Barcode-variant association was carried out with a tailor software based on MPRAflow. RNA to DNA ratio was also counted by our tailored version of MPRAflow [44]. MPRAnalyze was used to calculate an alpha factor that represents transcriptional activity [48]. Any variant not represented by at least 10 barcodes was excluded from the library (*n*_*variants*_ = 1, 301). The log_2_ fold change was calculated by taking the difference of the log_2_ of the alpha factor and the mean of the log_2_ of the control activity (equivalent to log_2_ of the geometric mean). The control activity was estimated from the alpha factors of selected variants from Run 18 (random sequences) and Generation 1 (1 mutation from wildtype) that had the lowest variance in their measured barcodes (the overlap of the 25% quantile in two replicates, n =33). The same calculation was used for computing the model predicted log_2_ fold change, but the predicted gene expression count was used in place of the alpha factor.

#### Cloning top variants

The top variants were synthesized by Twist Bioscience. Fragments were assembled into the cloning vector described above, without the degenerate barcode.

### Microscopy and Fluorescence Quantification

Imaging of variants was carried out 20-24 h after transfection. A Leica DM2000 upright microscope with K5 camera, LED excitation bulb and GFP filter for emission (520nm). Images were acquired with LASX software. All images were acquired with the same bulb intensity at an exposure time of 1 msec, Images were further processed in Fiji [78]. The background signal was removed from the GFP channel by measuring the mean gray value of a small patch between cells and subtracting this value plus two standard deviations for each individual pane of view. The maximum brightness is set to 35,000 on all images displayed. Fluorescent signal was quantified by using the threshold tool in FIJI followed by Analyze Particles. The raw integrated density for each particle was measured and divided by the area. Results were plotted in python using the package seaborn and statistical tests were performed in R (http://www.r-project.org).

### Fluorescence Activated Cell Sorting

Cells were sorted on a BD FACSAria sorter 24 hours after transfection at Queen Mary University London. Analysis of cell brightness was carried out in FlowJo software, with gating settings detailed in Fig.S4.

### Motif Analysis

A list of 319 defined motifs in tomato from Plant TFDB ([52] accessed at https://planttfdb.gao-lab.org/) was used to search (1) the wildtype sequence and (2) variant 3 using FIMO with a p-value cut off of *<* 1*e*^−5^ [51].

## Data Availability

Reference genome sequences and annotations were obtained from Ensembl Plants [61] release-56 (https://plants.ensembl.org/index.html).

Raw RNA-seq data was downloaded from National Center for Biotechnology Information, National Library of Medicine, National Institute of Health (NCBI NLM NIH, https://www.ncbi.nlm.nih.gov/). Pro-cessed RNA-seq data was downloaded from PlantExp database (https://biotec.njau.edu.cn/plantExp/). Processed ChIP-Seq, ATAC-seq, DNAse-seq and Faire-seq data was downloaded from ChIP Hub (https://biobigdata.nju.edu.cn/ChIPHub/). Plant Transcription Factor Binding Site Motifs were obtained from PlantTFDB (https://planttfdb.gao-lab.org/).

## Author’s Contributions

Conceptualisation: S.J. and N.K. Analysis: S.J., A.B., T.F., C.D., C.G., N.K. Writing: S.J., C.D., J.C., N.K.

## Acknowledgements

We thank Shaoli Das Gupta and the wider Phytoform Labs team for their contribution to CRE.AI.TIVE, Gary Warrnes from Queen Mary University of London for support with cell sorting, and Tara Advaney, Manuel Gomez, Kevin Shaffer-Morrison and Brian Skjerven from Amazon Web Services for their support with cloud computing.

## References

[1] Baenziger, P. S., Mumm, R. H., Bernardo, R., Brummer, C. E., Langridge, P., Simon, P., et al. “Plant Breeding and Genetics—A paper in the series on The Need for Agricultural Innovation to Sustainably Feed the World by 2050”. In: Council for Agricultural Science and Technology (CAST) (57 2017).

[2] Kafle, S. “CRISPR/CAS9: A new paradigm for crop improvement revolutionizing agriculture”. In: Journal of Agriculture and Food Research 11 (2023). issn: 26661543. doi: 10.1016/j.jafr.2022.100484.

[3] Hanika, K., Schipper, D., Chinnappa, S., Oortwijn, M., Schouten, H. J., Thomma, B. P., et al. “Impairment of Tomato WAT1 Enhances Resistance to Vascular Wilt Fungi Despite Severe Growth Defects”. In: Frontiers in Plant Science 12 (2021). issn: 1664462X. doi: 10.3389/fpls.2021.721674.

[4] Kodackattumannil, P., Lekshmi, G., Kottackal, M., Sasi, S., Krishnan, S., Senaani, S. A., et al. “Hidden pleiotropy of agronomic traits uncovered by CRISPR-Cas9 mutagenesis of the tyrosinase CuA-binding domain of the polyphenol oxidase 2 of eggplant”. In: Plant Cell Reports 42 (4 2023), pp. 825–828. issn: 1432203X. doi: 10.1007/s00299-023-02987-x.

[5] Vasudevan, S. N., Pooja, S. K., Raju, T. J., and Damini, C. S. “Cisgenics and intragenics: boon or bane for crop improvement”. In: Frontiers in Plant Science 14 (2023). issn: 1664462X. doi: 10.3389/fpls.2023.1275145.

[6] Rodríguez-Leal, D., Lemmon, Z. H., Man, J., Bartlett, M. E., and Lippman, Z. B. “Engineering Quantitative Trait Variation for Crop Improvement by Genome Editing”. In: Cell 171 (2 2017), 470–480.e8. issn: 10974172. doi: 10.1016/j.cell.2017.08.030.

[7] Hendelman, A., Zebell, S., Rodriguez-Leal, D., Dukler, N., Robitaille, G., Wu, X., et al. “Conserved pleiotropy of an ancient plant homeobox gene uncovered by cis-regulatory dissection”. In: Cell 184 (7 2021), 1724–1739.e16. issn: 10974172. doi: 10.1016/j.cell.2021.02.001.

[8] Wittkopp, P. J. and Kalay, G. “Cis-regulatory elements: molecular mechanisms and evolutionary processes underlying divergence”. In: Nature Reviews Genetics 13 (1 2012), pp. 59–69. issn: 1471-0064. doi: 10.1038/nrg3095. url: https://doi.org/10.1038/nrg3095.

[9] Zhou, J., Liu, G., Zhao, Y., Zhang, R., Tang, X., Li, L., et al. “An efficient CRISPR–Cas12a promoter editing system for crop improvement”. In: Nature Plants 9 (4 2023), pp. 588–604. issn: 20550278. doi: 10.1038/s41477-023-01384-2.

[10] Liu, L., Gallagher, J., Arevalo, E. D., Chen, R., Skopelitis, T., Wu, Q., et al. “Enhancing grain-yield-related traits by CRISPR–Cas9 promoter editing of maize CLE genes”. In: Nature Plants 7 (3 2021), pp. 287–294. issn: 20550278. doi: 10.1038/s41477-021-00858-5.

[11] Wang, X., Aguirre, L., Rodríguez-Leal, D., Hendelman, A., Benoit, M., and Lippman, Z. B. “Dissecting cis-regulatory control of quantitative trait variation in a plant stem cell circuit”. In: Nature Plants 7 (4 2021), pp. 419–427. issn: 20550278. doi: 10.1038/s41477-021-00898-x.

[12] Song, X., Meng, X., Guo, H., Cheng, Q., Jing, Y., Chen, M., et al. “Targeting a gene regulatory element enhances rice grain yield by decoupling panicle number and size”. In: Nature Biotechnology 40 (9 2022), pp. 1403–1411. issn: 15461696. doi: 10.1038/s41587-022-01281-7.

[13] Bhunia, R. K., Menard, G. N., and Eastmond, P. J. “A native promoter–gene fusion created by CRISPR/Cas9-mediated genomic deletion offers a transgene-free method to drive oil accumulation in leaves”. In: FEBS Letters 596 (15 2022), pp. 1865–1870. issn: 18733468. doi: 10.1002/1873-3468.14365.

[14] Patel-Tupper, D., Kelikian, A., Leipertz, A., Maryn, N., Tjahjadi, M., Karavolias, N. G., et al. “Multiplexed CRISPR-Cas9 mutagenesis of rice PSBS1 noncoding sequences for transgene-free overexpression”. In: Sci. Adv 10 (2024), p. 7452. url: https://www.science.org.

[15] Dong, O. X. and Ronald, P. C. “Targeted DNA insertion in plants”. In: Proceedings of the National Academy of Sciences 118 (22 2021), pp. 1–9. doi: 10.1073/pnas.2004834117/-/DCSupplemental. url: https://www.pnas.org/page/collection/.

[16] Gao, H., Mutti, J., Young, J. K., Yang, M., Schroder, M., Lenderts, B., et al. “Complex Trait Loci in Maize Enabled by CRISPR-Cas9 Mediated Gene Insertion”. In: Frontiers in Plant Science 11 (2020). issn: 1664462X. doi: 10.3389/fpls.2020.00535.

[17] Lu, Y., Tian, Y., Shen, R., Yao, Q., Wang, M., Chen, M., et al. “Targeted, efficient sequence insertion and replacement in rice”. In: Nature Biotechnology 38 (12 2020), pp. 1402–1407. issn: 1087-0156, 1546-1696. doi: 10.1038/s41587-020-0581-5. url: https://www.nature.com/articles/s41587-020-0581-5files/116/Luetal.-2020-Targeted,efficientsequenceinsertionandreplace.pdf.

[18] Yao, Q., Shen, R., Shao, Y., Tian, Y., Han, P., Zhang, X., et al. “Efficient and multiplex gene upregulation in plants through CRISPR-Cas-mediated knockin of enhancers”. In: Molecular plant 17 (9 2024), pp. 1472–1483. issn: 17529867. doi: 10.1016/j.molp.2024.07.009.

[19] Jores, T., Mueth, N. A., Tonnies, J., Char, S. N., Liu, B., Grillo-Alvarado, V., et al. “Small DNA elements that act as both insulators and silencers in plants”. In: BioRxiv (2024). doi: 10.1101/2024.09.13.612883. url: http://biorxiv.org/lookup/doi/10.1101/2024.09.13.612883.

[20] Akagi, T., Masuda, K., Kuwada, E., Takeshita, K., Kawakatsu, T., Ariizumi, T., et al. “Genome-wide cis-decoding for expression design in tomato using cistrome data and explainable deep learning”. In: Plant Cell 34 (6 2022), pp. 2174–2187. issn: 1532298X. doi: 10.1093/plcell/koac079.

[21] Washburn, J. D., Mejia-Guerra, M. K., Ramstein, G., Kremling, K. A., Valluru, R., Buckler, E. S., et al. “Evolutionarily informed deep learning methods for predicting relative transcript abundance from DNA sequence”. In: Proceedings of the National Academy of Sciences of the United States of America 116 (12 2019), pp. 5542–5549. issn: 10916490. doi: 10.1073/pnas.1814551116.

[22] Peleke, F. F., Zumkeller, S. M., Gültas, M., Schmitt, A., and Szymański, J. “Deep learning the cis-regulatory code for gene expression in selected model plants”. In: Nature Communications 15 (1 2024), p. 3488. issn: 2041-1723. doi: 10.1038/s41467-024-47744-0. url: https://www.nature.com/articles/s41467-024-47744-0.

[23] Mendoza-Revilla, J., Trop, E., Gonzalez, L., Roller, M., Dalla-Torre, H., Almeida B. P. de, et al. “A foundational large language model for edible plant genomes”. In: Communications Biology 7 (1 2024). issn: 23993642. doi: 10.1038/s42003-024-06465-2.

[24] Avsec, iga, Agarwal, V., Visentin, D., Ledsam, J. R., Grabska-Barwinska, A., Taylor, K. R., et al. “Effective gene expression prediction from sequence by integrating long-range interactions”. In: Nature methods (2021). doi: 10.1038/s41592-021-01252-x. url: https://doi.org/10.1038/s41592-021-01252-x.

[25] Linder, J., Srivastava, D., Yuan, H., Agarwal, V., and Kelley, D. R. “Predicting RNA-seq coverage from DNA sequence as a unifying model of gene regulation”. In: BioRxiv (2023). doi: 10.1101/2023.08.30.555582. url: http://biorxiv.org/lookup/doi/10.1101/2023.08.30.555582.

[26] Ji, Y., Zhou, Z., Liu, H., and Davuluri, R. V. “DNABERT: pre-trained Bidirectional Encoder Representations from Transformers model for DNA-language in genome”. In: Bioinformatics 37 (15 2021), pp. 2112–2120. issn: 13674811. doi: 10.1093/bioinformatics/btab083.

[27] Dalla-Torre, H., Gonzalez, L., Mendoza-Revilla, J., Carranza, N. L., Grzywaczewski, A. H., Oteri, F., et al. “Nucleotide Transformer: building and evaluating robust foundation models for human genomics”. In: Nature Methods (2024). issn: 1548-7091. doi: 10.1038/s41592-024-02523-z. url: https://www.nature.com/articles/s41592-024-02523-z.

[28] Nguyen, E., Poli, M., Durrant, M. G., Kang, B., Katrekar, D., Li, D. B., et al. “Sequence modeling and design from molecular to genome scale with Evo”. In: Science 386 (6723 2024). issn: 0036-8075. doi: 10.1126/science.ado9336. url: https://www.science.org/doi/10.1126/science.ado9336.

[29] Saberi, A., Choi, B., Wang, S., Hernández-Corchado, A., Naghipourfar, M., Namini, A. M., et al. “A long-context RNA foundation model for predicting transcriptome architecture”. In: BioRxiv (2024). doi: 10.1101/2024.08.26.609813. url: http://biorxiv.org/lookup/doi/10.1101/2024.08.26.609813.

[30] Benegas, G. I., Batra, S. I. S., Song, Y. S., and Kathryn Roeder, I. E. by. “DNA language models are powerful predictors of genome-wide variant effects”. In: Proceedings of the National Academy of Sciences 120 (44 2023), pp. 1–9. doi: 10.1073/pnas.

[31] Schiff, Y., Kao, C.-H., Gokaslan, A., Dao, T., Gu, A., and Kuleshov, V. “Caduceus: Bi-Directional Equivariant Long-Range DNA Sequence Modeling”. In: Arxiv (2024). url: http://arxiv.org/abs/2403.03234.

[32] Nguyen, E., Poli, M., Faizi, M., Thomas, A., Birch-Sykes, C., Wornow, M., et al. “HyenaDNA: Long-Range Genomic Sequence Modeling at Single Nucleotide Resolution”. In: Arxiv (2023). url: http://arxiv.org/abs/2306.15794.

[33] Zhou, J. and Troyanskaya, O. G. “Predicting effects of noncoding variants with deep learning-based sequence model”. In: Nature Methods 12 (10 2015), pp. 931–934. issn: 15487105. doi: 10.1038/nmeth.3547.

[34] Chen, K. M., Wong, A. K., Troyanskaya, O. G., and Zhou, J. “A sequence-based global map of regulatory activity for deciphering human genetics”. In: Nature Genetics 54 (7 2022), pp. 940–949. issn: 15461718. doi: 10.1038/s41588-022-01102-2.

[35] Zhou, J., Theesfeld, C. L., Yao, K., Chen, K. M., Wong, A. K., and Troyanskaya, O. G. “Deep learning sequence-based ab initio prediction of variant effects on expression and disease risk”. In: Nature Genetics 50 (8 2018), pp. 1171–1179. issn: 15461718. doi: 10.1038/s41588-018-0160-6.

[36] Agarwal, V., Inoue, F., Schubach, M., Martin, B. K., Dash, P. M., Zhang, Z., et al. “Massively parallel characterization of transcriptional regulatory elements in three diverse human cell types”. In: BioRxiv (2023). doi: 10.1101/2023.03.05.531189. url: http://biorxiv.org/lookup/doi/10.1101/2023.03.05.531189.

[37] Kelley, D. R., Reshef, Y. A., Bileschi, M., Belanger, D., McLean, C. Y., and Snoek, J. “Sequential regulatory activity prediction across chromosomes with convolutional neural networks”. In: Genome Research 28 (5 2018), pp. 739–750. issn: 15495469. doi: 10.1101/gr.227819.117.

[38] Kelley, D. R. “Cross-species regulatory sequence activity prediction”. In: PLoS Computational Biology 16 (7 2020). issn: 15537358. doi: 10.1371/journal.pcbi.1008050.

[39] Agarwal, V. and Shendure, J. “Predicting mRNA Abundance Directly from Genomic Sequence Using Deep Convolutional Neural Networks”. In: Cell Reports 31 (7 2020). issn: 22111247. doi: 10.1016/j.celrep.2020.107663.

[40] Toneyan, S. and Koo, P. K. “Interpreting cis-regulatory interactions from large-scale deep neural networks”. In: Nature Genetics 56 (11 2024), pp. 2517–2527. issn: 1546-1718. doi: 10.1038/s41588-024-01923-3. url: https://doi.org/10.1038/s41588-024-01923-3.

[41] Devlin, J., Chang, M.-W., Lee, K., Google, K. T., and Language, A. I. BERT: Pre-training of Deep Bidirectional Transformers for Language Understanding. url: https://github.com/tensorflow/tensor2tensor.

[42] Liang, Y., Ma, F., Li, B., Guo, C., Hu, T., Zhang, M., et al. “A bHLH transcription factor, SlbHLH96, promotes drought tolerance in tomato”. In: Horticulture Research 9 (2022). issn: 20527276. doi: 10.1093/hr/uhac198.

[43] Jores, T., Tonnies, J., Wrightsman, T., Buckler, E. S., Cuperus, J. T., Fields, S., et al. “Synthetic promoter designs enabled by a comprehensive analysis of plant core promoters”. In: Nature Plants 7 (6 2021), pp. 842–855. issn: 20550278. doi: 10.1038/s41477-021-00932-y.

[44] Gordon, M. G., Inoue, F., Martin, B., Schubach, M., Agarwal, V., Whalen, S., et al. “lentiMPRA and MPRAflow for high-throughput functional characterization of gene regulatory elements”. In: Nature Protocols 15 (8 2020), pp. 2387–2412. issn: 17502799. doi: 10.1038/s41596-020-0333-5.

[45] Kim, J. Y., Symeonidi, E., Pang, T. Y., Denyer, T., Weidauer, D., Bezrutczyk, M., et al. “Distinct identities of leaf phloem cells revealed by single cell transcriptomics”. In: Plant Cell 33 (3 2021), pp. 511–530. issn: 1532298X. doi: 10.1093/plcell/koaa060.

[46] Tiwari, P., Indoliya, Y., Chauhan, A. S., Singh, P., Singh, P. K., Singh, P. C., et al. “Auxin-salicylic acid cross-talk ameliorates OsMYB–R1 mediated defense towards heavy metal, drought and fungal stress”. In: Journal of Hazardous Materials 399 (2020). issn: 18733336. doi: 10.1016/j.jhazmat.2020.122811.

[47] Bolduc, N., Yilmaz, A., Mejia-Guerra, M. K., Morohashi, K., O’Connor, D., Grotewold, E., et al. “Unraveling the KNOTTED1 regulatory network in maize meristems”. In: Genes and Development 26 (15 2012), pp. 1685–1690. issn: 08909369. doi: 10.1101/gad.193433.112.

[48] Ashuach, T., Fischer, D. S., Kreimer, A., Ahituv, N., Theis, F. J., and Yosef, N. “MPRAnalyze: Statistical framework for massively parallel reporter assays”. In: Genome Biology 20 (1 2019). issn: 1474760X. doi: 10.1186/s13059-019-1787-z.

[49] Boer, D. R., Freire-Rios, A., Berg, W. A. V. D., Saaki, T., Manfield, I. W., Kepinski, S., et al. “Structural basis for DNA binding specificity by the auxin-dependent ARF transcription factors”. In: Cell 156 (3 2014), pp. 577–589. issn: 10974172. doi: 10.1016/j.cell.2013.12.027.

[50] Adams, C. C. and Workman, J. L. “Binding of disparate transcriptional activators to nucleosomal DNA is inherently cooperative”. In: Molecular and Cellular Biology 15 (3 1995). doi: 10.1128/MCB.15.3.1405, pp. 1405–1421. issn: null. DOI: 10.1128/MCB.15.3.1405. url: https://doi.org/10.1128/MCB.15.3.1405.

[51] Grant, C. E., Bailey, T. L., and Noble, W. S. “FIMO: Scanning for occurrences of a given motif”. In: Bioinformatics 27 (7 2011), pp. 1017–1018. issn: 14602059. doi: 10.1093/bioinformatics/btr064.

[52] Jin, J., Tian, F., Yang, D. C., Meng, Y. Q., Kong, L., Luo, J., et al. “PlantTFDB 4.0: Toward a central hub for transcription factors and regulatory interactions in plants”. In: Nucleic Acids Research 45 (D1 2017), pp. D1040–D1045. issn: 13624962. doi: 10.1093/nar/gkw982.

[53] Jumper, J., Evans, R., Pritzel, A., Green, T., Figurnov, M., Ronneberger, O., et al. “Highly accurate protein structure prediction with AlphaFold”. In: Nature 596 (7873 2021), pp. 583–589. issn: 14764687. doi: 10.1038/s41586-021-03819-2.

[54] Benfey, P. N. and Chua N. hai. “The Cauliflower Mosaic Virus 35S Promoter: Combinatorial Regulation of Transcription in Plants”. In: Science 161 (1985), p. 2024. url: https://www.science.org.

[55] Liu, Q., Kasuga, M., Sakuma, Y., Abe, H., Miura, S., Yamaguchi-Shinozaki, K., et al. “Two Transcription Factors, DREB1 and DREB2, with an EREBP/AP2 DNA Binding Domain Separate Two Cellular Signal Transduction Pathways in Drought- and Low-Temperature-Responsive Gene Expression, Respectively, in Arabidopsis”. In: The Plant Cell 10 (8 1998), pp. 1391–1406. issn: 1040-4651. doi: 10.1105/tpc.10.8.1391. url: https://doi.org/10.1105/tpc.10.8.1391.

[56] Cai, Y. M., Kallam, K., Tidd, H., Gendarini, G., Salzman, A., and Patron, N. J. “Rational design of minimal synthetic promoters for plants”. In: Nucleic Acids Research 48 (21 2021), pp. 11845–11856. issn: 13624962. doi: 10.1093/nar/gkaa682.

[57] Cai, Y. M., Witham, S., and Patron, N. J. “Tuning Plant Promoters Using a Simple Split Luciferase Method to Assess Transcription Factor-DNA Interactions”. In: ACS Synthetic Biology 12 (11 2023), pp. 3482–3486. issn: 21615063. doi: 10.1021/acssynbio.3c00094.

[58] Voichek, Y., Hristova, G., Mollá-Morales, A., Weigel, D., and Nordborg, M. “Widespread position-dependent transcriptional regulatory sequences in plants”. In: Nature Genetics (2024). issn: 15461718. doi: 10.1038/s41588-024-01907-3.

[59] Schöne, S., Jurk, M., Helabad, M. B., Dror, I., Lebars, I., Kieffer, B., et al. “Sequences flanking the core-binding site modulate glucocorticoid receptor structure and activity”. In: Nature Communications 7 (2016). issn: 20411723. doi: 10.1038/ncomms12621.

[60] Karollus, A., Mauermeier, T., and Gagneur, J. “Current sequence-based models capture gene expression determinants in promoters but mostly ignore distal enhancers”. In: Genome Biology 24 (1 2023). issn: 1474760X. doi: 10.1186/s13059-023-02899-9.

[61] Harrison, P. W., Amode, M. R., Austine-Orimoloye, O., Azov, A. G., Barba, M., Barnes, I., et al. “Ensembl 2024”. In: Nucleic Acids Research 52 (D1 2024), pp. D891–D899. issn: 13624962. doi: 10.1093/nar/gkad1049.

[62] Fu, L. Y., Zhu, T., Zhou, X., Yu, R., He, Z., Zhang, P., et al. “ChIP-Hub provides an integrative platform for exploring plant regulome”. In: Nature Communications 13 (1 2022). issn: 20411723. doi: 10.1038/s41467-022-30770-1.

[63] Liu, J., Zhang, Y., Zheng, Y., Zhu, Y., Shi, Y., Guan, Z., et al. “PlantExp: a platform for exploration of gene expression and alternative splicing based on public plant RNA-seq samples”. In: Nucleic Acids Research 51 (1 D 2023), pp. D1483–D1491. issn: 13624962. doi: 10.1093/nar/gkac917.

[64] NIH NCBI. Sequence Read Archive (SRA) on AWS. https://registry.opendata.aws/ncbi-sra. Accessed on 9th May 2024 and 11th September 2024.

[65] Ewels, P. A., Peltzer, A., Fillinger, S., Patel, H., Alneberg, J., Wilm, A., et al. “The nf-core framework for community-curated bioinformatics pipelines”. In: Nature Biotechnology 38 (3 2020), pp. 272–276. issn: 15461696. doi: 10.1038/s41587-020-0439-x.

[66] Kim, D., Paggi, J. M., Park, C., Bennett, C., and Salzberg, S. L. “Graph-based genome alignment and genotyping with HISAT2 and HISAT-genotype”. In: Nature Biotechnology 37 (8 2019), pp. 907–915. issn: 15461696. doi: 10.1038/s41587-019-0201-4.

[67] Ramírez, F., Ryan, D. P., Grüning, B., Bhardwaj, V., Kilpert, F., Richter, A. S., et al. “deepTools2: a next generation web server for deep-sequencing data analysis”. In: Nucleic Acids Research 44 (W1 2016), W160–W165. issn: 13624962. doi: 10.1093/NAR/GKW257.

[68] Li, H. “Aligning sequence reads, clone sequences and assembly contigs with BWA-MEM”. In: Arxiv (2013). url: http://arxiv.org/abs/1303.3997.

[69] Li, H., Handsaker, B., Wysoker, A., Fennell, T., Ruan, J., Homer, N., et al. “The Sequence Alignment/Map format and SAMtools”. In: Bioinformatics 25 (16 2009), pp. 2078–2079. issn: 13674803. doi: 10.1093/bioinformatics/btp352.

[70] Pertea, M., Pertea, G. M., Antonescu, C. M., Chang, T. C., Mendell, J. T., and Salzberg, S. L. “StringTie enables improved reconstruction of a transcriptome from RNA-seq reads”. In: Nature Biotechnology 33 (3 2015), pp. 290–295. issn: 15461696. doi: 10.1038/nbt.3122.

[71] Pertea, M., Kim, D., Pertea, G. M., Leek, J. T., and Salzberg, S. L. “Transcript-level expression analysis of RNA-seq experiments with HISAT, StringTie and Ballgown”. In: Nature Protocols 11 (9 2016), pp. 1650–1667. issn: 1750-2799. doi: 10.1038/nprot.2016.095. url: https://doi.org/10.1038/nprot.2016.095.

[72] Kovaka, S., Zimin, A. V., Pertea, G. M., Razaghi, R., Salzberg, S. L., and Pertea, M. “Transcriptome assembly from long-read RNA-seq alignments with StringTie2”. In: Genome Biology 20 (1 2019). issn: 1474760X. doi: 10.1186/s13059-019-1910-1.

[73] Shumate, A., Wong, B., Pertea, G., and Pertea, M. “Improved transcriptome assembly using a hybrid of long and short reads with StringTie”. In: PLoS Computational Biology 18 (6 2022). issn: 15537358. doi: 10.1371/journal.pcbi.1009730.

[74] Zhou, Y., Zhang, Z., Bao, Z., Li, H., Lyu, Y., Zan, Y., et al. “Graph pangenome captures missing heritability and empowers tomato breeding”. In: Nature 606 (7914 2022), pp. 527–534. issn: 14764687. doi: 10.1038/s41586-022-04808-9.

[75] Fernandez-Pozo, N., Menda, N., Edwards, J. D., Saha, S., Tecle, I. Y., Strickler, S. R., et al. “The Sol Genomics Network (SGN)-from genotype to phenotype to breeding”. In: Nucleic Acids Research 43 (D1 2015), pp. D1036–D1041. issn: 13624962. doi: 10.1093/nar/gku1195.

[76] Lampropoulos, A, Sutikovic, Z, Wenzl, C, Maegele, I, and Lohmann, J. U. “GreenGate-A Novel, Versatile, and Efficient Cloning System for Plant Transgenesis”. In: PLoS ONE 8 (12 2013), p. 83043. doi: 10.1371/journal.pone.0083043. url: www.plosone.org.

[77] Nicolia, A., Fält, A.-S., Hofvander, P., and Andersson, M. “Protoplast-Based Method for Genome Editing in Tetraploid Potato”. In: ed. by P. Tripodi. Springer US, 2021, pp. 177–186. isbn: 978-1-0716-1201-9. doi: 10.1007/978-1-0716-1201-9_12. url: https://doi.org/10.1007/978-1-0716-1201-9_12.

[78] Schindelin, J., Arganda-Carreras, I., Frise, E., Kaynig, V., Longair, M., Pietzsch, T., et al. “Fiji: an open-source platform for biological-image analysis”. In: Nature Methods 9 (7 2012), pp. 676–682. issn: 1548-7105. doi: 10.1038/nmeth.2019. url: https://doi.org/10.1038/nmeth.2019.

