## Supplementary Material for "A scalable method for modulating plant gene expression using a multispecies genomic model and protoplast-based massively parallel reporter assay"

### Supplementary Materials

Table S1: The plant species and number of transcriptomic and epigenomic dataset used to pre-train the CRE.AI.TIVE model.

Table S2: Summary of FIMO-identified motifs on variant 3 and wild-type proximal promoter sequence.

Figure S1: Examples of RNA-seq coverage across all species.

Figure S2: Examples of scatter plots of predicted vs true gene expression $\log_2$(count) values for an individual species and an individual experimental condition.

Figure S3: Examples of Scatter plots of predicted vs true gene expression $\log_2$(count) values for an individual gene in an individual species across all experimental RNA-seq conditions.

Figure S4: Gating boundaries for FACS-sorted cells.

Supplementary references.

**Table S1 - The plant species and number of transcriptomic and epigenomic dataset used to pre-train the CRE.AI.TIVE model**

| Species | Experiment | Num samples | Num targets |
|---|---|---|---|
| Arabidopsis Lyrata | RNA-seq | 8 | 5 |
| | ChIP-seq | 10 | 5 |
| Arabidopsis Thaliana | RNA-seq | 3595 | 1499 |
| | ATAC-seq | 53 | 53 |
| | ChIP-seq | 1344 | 542 |
| | DNase-seq | 14 | 14 |
| | FAIRE-seq | 1 | 1 |
| Brachypodium Distachyon | RNA-seq | 102 | 73 |
| | ATAC-seq | 3 | 3 |
| | ChIP-seq | 2 | 1 |
| | DNase-seq | 18 | 18 |
| Citrullus Lanatus | RNA-seq | 6 | 2 |
| Cucumis Sativus | RNA-seq | 7 | 2 |
| | ChIP-seq | 27 | 12 |
| | DNase-seq | 10 | 10 |
| Glycine Max | RNA-seq | 51 | 17 |
| | ATAC-seq | 12 | 12 |
| | ChIP-seq | 96 | 27 |
| Medicago Truncatula | RNA-seq | 204 | 60 |
| | ATAC-seq | 9 | 9 |
| | ChIP-seq | 24 | 8 |
| Oryza Sativa | RNA-seq | 236 | 93 |
| | ATAC-seq | 5 | 5 |
| | ChIP-seq | 162 | 105 |
| | DAP-seq | 2 | 2 |
| | DNase-seq | 2 | 2 |
| Setaria Italica | RNA-seq | 16 | 8 |
| | ChIP-seq | 4 | 3 |
| | DNase-seq | 18 | 18 |
| Solanum Lycopersicum | RNA-seq | 174 | 53 |
| Triticum Aestivum | RNA-seq | 31 | 14 |
| | ATAC-seq | 1 | 1 |
| | ChIP-seq | 20 | 14 |
| | DNase-seq | 3 | 3 |
| Zea Mays | RNA-seq | 488 | 150 |

**Table S2 - Summary of FIMO-identified motifs on variant 3 and wild-type proximal promoter sequence.**

| binding gene id | bound variant | motif sequence 5' - 3' | strand | p-value | transcription factor family | 5' distance from TSS | 3' distance from TSS |
|---|---|---|---|---|---|---|---|
| Solyc04g008870.2 | variant 3 | TGTGTGGATAAGG | forward | 3.57E-06 | MYB | 219 | 207 |
| Solyc07g026680.2 | variant 3 | AGCCTTATCCACACA | reverse | 4.46E-06 | MYB | 219 | 205 |
| Solyc06g071230.2 | variant 3 | GTGTGGATAAGGCT | forward | 3.75E-06 | MYB | 218 | 205 |
| Solyc08g078340.2 | variant 3 | TAAGCCTTATCCACA | reverse | 7.50E-07 | MYB | 217 | 203 |
| Solyc03g119740.2 | variant 3 | ATAAGCCTTATCCAC | reverse | 8.90E-06 | MYB | 216 | 202 |
| Solyc04g005100.2 | variant 3 | GTGGATAAGG | forward | 4.03E-06 | MYB | 216 | 207 |
| Solyc05g055240.2 | variant 3 | TGGATAAGGC | forward | 8.14E-06 | MYB-like | 215 | 206 |
| Solyc11g006720.1 | variant 3 | TGGATAAGGC | forward | 8.14E-06 | MYB | 215 | 206 |
| Solyc04g008870.2 | variant 3 | TGTGTGGATAAGG | forward | 3.57E-06 | MYB | 190 | 178 |
| Solyc07g026680.2 | variant 3 | AACCTTATCCACACA | reverse | 5.98E-07 | MYB | 190 | 176 |
| Solyc03g096350.2 | variant 3 | AACCTTATCCACAC | reverse | 2.03E-06 | MYB | 189 | 176 |
| Solyc06g071230.2 | variant 3 | GTGTGGATAAGGTT | forward | 8.06E-07 | MYB | 189 | 176 |
| Solyc08g078340.2 | variant 3 | GCAACCTTATCCACA | reverse | 1.76E-06 | MYB | 188 | 174 |
| Solyc04g005100.2 | variant 3 | GTGGATAAGG | forward | 4.03E-06 | MYB | 187 | 178 |
| Solyc05g055240.2 | variant 3 | TGGATAAGGT | forward | 6.12E-06 | MYB-like | 186 | 177 |
| Solyc11g006720.1 | variant 3 | TGGATAAGGT | forward | 6.12E-06 | MYB | 186 | 177 |
| Solyc04g078840.2 | variant 3 | AAGGTTGCCACGTGTAAC | forward | 8.67E-06 | Bzip | 182 | 165 |
| Solyc08g005290.2 | variant 3 | TGGTTACACGTGGCA | reverse | 6.42E-06 | Bzip | 177 | 163 |
| Solyc10g081350.1 | variant 3 | TGGTTACACGTGGCA | reverse | 8.35E-06 | Bzip | 177 | 163 |
| Solyc09g014250.2 | variant 3 | GATTTTGTGGATAAGGTTGGT | reverse | 1.26E-06 | MYB | 166 | 146 |
| Solyc05g055240.2 | variant 3 | TGGATAAGGT | reverse | 6.12E-06 | MYB-like | 161 | 152 |
| Solyc11g006720.1 | variant 3 | TGGATAAGGT | reverse | 6.12E-06 | MYB | 161 | 152 |
| Solyc04g005100.2 | variant 3 | GTGGATAAGG | reverse | 4.03E-06 | MYB | 160 | 151 |
| Solyc09g014250.2 | variant 3 | AGAGGATAAGATAAGATTTTG | reverse | 9.47E-06 | MYB | 152 | 132 |
| Solyc03g096350.2 | variant 3 | TATCTTATCCTCTT | forward | 5.11E-06 | MYB | 143 | 130 |
| Solyc04g008870.2 | variant 3 | TAAGAAGATAAGA | forward | 9.47E-06 | MYB | 99 | 87 |
| Solyc03g113760.2 | wildtype | GGTGAAGTCTTCCCGCAAATA | forward | 7.69E-06 | E2F | 184 | 163 |
| Solyc06g076030.2 | wildtype | TTTATTTTTTATTATATTTTT | reverse | 1.46E-06 | Dof zinc finger protein | 95 | 74 |
| Solyc06g076030.2 | wildtype | TTAGTTTTTTATTTTTTATTA | reverse | 3.91E-06 | Dof zinc finger protein | 88 | 67 |
| Solyc11g008560.1 | wildtype | AATAAAAAATAAAAAACTAA | forward | 2.21E-06 | ERF | 87 | 67 |

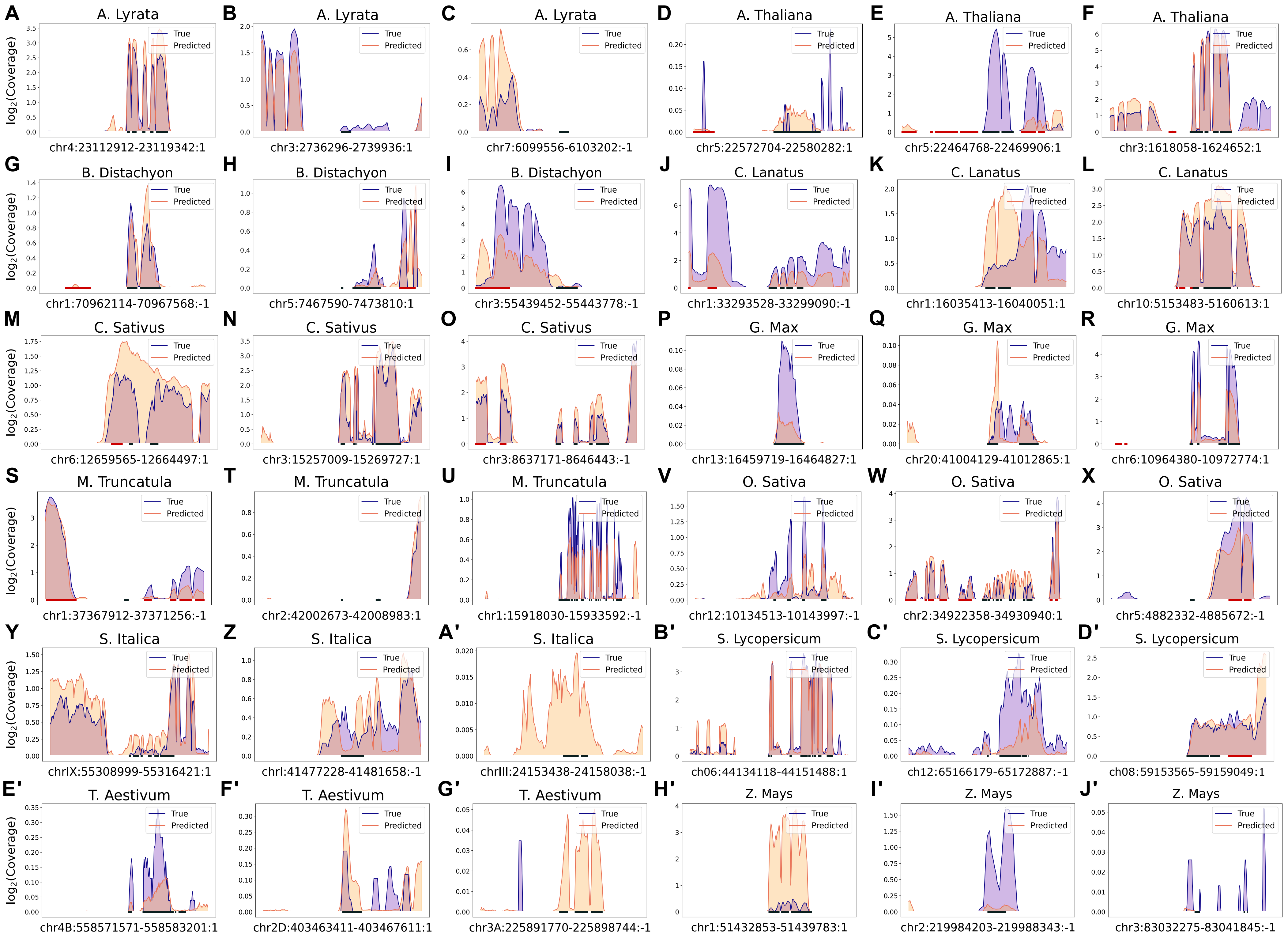

log$_2$(Coverage)
True
Predicted
A
A. Lyrata
chr4:23112912-23119342:1
B
A. Lyrata
chr3:2736296-2739936:1
C
A. Lyrata
chr7:6099556-6103202:-1
D
A. Thaliana
chr5:22572704-22580282:1
E
A. Thaliana
chr5:22464768-22469906:1
F
A. Thaliana
chr3:1618058-1624652:1
G
B. Distachyon
chr1:70962114-70967568:-1
H
B. Distachyon
chr5:7467590-7473810:1
I
B. Distachyon
chr3:55439452-55443778:-1
J
C. Lanatus
chr1:33293528-33299090:-1
K
C. Lanatus
chr1:16035413-16040051:1
L
C. Lanatus
chr10:5153483-5160613:1
M
C. Sativus
chr6:12659565-12664497:1
N
C. Sativus
chr3:15257009-15269727:1
O
C. Sativus
chr3:8637171-8646443:-1
P
G. Max
chr13:16459719-16464827:1
Q
G. Max
chr20:41004129-41012865:1
R
G. Max
chr6:10964380-10972774:1
S
M. Truncatula
chr1:37367912-37371256:-1
T
M. Truncatula
chr2:42002673-42008983:1
U
M. Truncatula
chr1:15918030-15933592:-1
V
O. Sativa
chr12:10134513-10143997:-1
W
O. Sativa
chr2:34922358-34930940:1
X
O. Sativa
chr5:4882332-4885672:-1
Y
S. Italica
chrIX:55308999-55316421:1
Z
S. Italica
chrI:41477228-41481658:-1
A'
S. Italica
chrIII:24153438-24158038:-1
B'
S. Lycopersicum
ch06:44134118-44151488:1
C'
S. Lycopersicum
ch12:65166179-65172887:-1
D'
S. Lycopersicum
ch08:59153565-59159049:1
E'
T. Aestivum
chr4B:558571571-558583201:1
F'
T. Aestivum
chr2D:403463411-403467611:1
G'
T. Aestivum
chr3A:225891770-225898744:-1
H'
Z. Mays
chr1:51432853-51439783:1
I'
Z. Mays
chr2:219984203-219988343:-1
J'
Z. Mays
chr3:83032275-83041845:-1

**Figure S1 - Examples of RNA-seq coverage across all species True (purple) Orange (predicted)**. **(A)** *Arabidopsis lyrata* gene fgenesh2_kg.4__2998__AT2G47580.1, pistil tissue 3 days after being infected with fusarium graminearum [1] **(B)** *Arabidopsis lyrata* gene scaffold_300762.1, mature cauline leaf [1] **(C)** *Arabidopsis lyrata* gene scaffold_701504.1, mature cauline leaf [1] **(D)** *Arabidopsis thaliana* gene AT5G55780, 5-6 week old male meiocyte [2] **(E)** *Arabidopsis thaliana* gene AT5G55450, 48 day old leaf tissue uncovered [3] **(F)** *Arabidopsis thaliana* gene AT3G05590, 7 day old seedling grown at 21 degrees under cool white light [4] **(G)** *Brachypodium distachyon* gene BRADI_1g73140v3, tissue from an emerging 3rd leaf 13 days after germination, collected after grown in continuous light and continuous 28 °C temperature for 15.5 hours [5] **(H)** *Brachypodium distachyon* gene BRADI_5g05832v3, tissue from an emerging 3rd leaf 13 days after germination, grown in continuous light and 12 hours per day of 28 °C temperature and 12 hours per day of 12 °C temperature [5] **(I)** *Brachypodium distachyon* gene BRADI_3g55436v3, tissue from an emerging 3rd leaf 13 days after germination, grown at continuous 28 °C temperature and 12 hours per day of light and 12 hours per day of dark [5] **(J)** *Citrullus lanatus* gene Cla97C01G020780, young-ripening fruit 30 days after pollination [6] **(K)** *Citrullus lanatus* gene Cla97C01G010510, young-ripening fruit 30 days after pollination [6] **(L)** *Citrullus lanatus* gene Cla97C10G189070, young fruit to ripening fruit 30 days after pollination [7] **(M)** Cucumis sativus gene Csa_6G197217, 1 month old tendril tissue [8] **(N)** *Cucumis sativus* gene Csa_3G238090, fruit 0 days after anthesis [9] **(O)** Cucumis sativus 1 month old tendril tissue [10] **(P)** *Glycine max* gene GLYMA_13G065300, embryo 45 days post initial seed (3-4mm) filling [11-15] **(Q)** *Glycine max* gene GLYMA_20G172400, embryo 55 days post initial seed (3-4mm) filling [11-15] **(R)** *Glycine max* gene GLYMA_06G133300, embryo 30 days post initial seed (3-4mm) filling [11-15] **(S)** *Medicago truncatula* gene MTR_1g083905, root 7 days after *Rhizoctonia solani* AG8 inoculation [16] **(T)** *Medicago truncatula* gene MTR_2g098270, root 2 days after Sinorhizobium medicae inoculation and treatment with limited nitrogen [17] **(U)** *Medicago truncatula* gene MTR_1g442790, shoot 18 days after planting, untreated [18] **(V)** *Oryza sativa* gene Os12g0275400, leaf tissue at the 4 leaf stage [19] **(W)** *Oryza sativa* gene Os02g0815400, shoot apical meristem [20,21] **(X)** *Oryza sativa* gene ENSRNA049450453, 3 year old callus tissue [22] **(Y)** *Setaria italica* gene SETIT_036985mg, L1 tiller bud 12 days after planting [23] **(Z)** *Setaria italica* gene SETIT_018931mg, 3 week old aboveground tissue [24] **(A')** *Setaria italica* gene SETIT_024937mg, 3 week old aboveground tissue [25] **(B')** *Solanum lycopersicum* gene Solyc06g074980.3, mature leaf tissue [26] **(C')** *Solanum lycopersicum* gene Solyc12g096940.3, generative cell tissue [27] **(D')** *Solanum lycopersicum* gene Solyc08g077120.2, mature root cell treated with phosphate deficiency for 24 hours [28,29] **(E')** *Triticum aestivum* gene TraesCS4B02G276900, seedling at the 3 leaf stage treated with ABA for 6 hours [30] **(F')** *Triticum aestivum* gene TraesCS2D02G313600, 2 month old root control (water) treatment [31] **(G')** *Triticum aestivum* gene TraesCS3A02G187800, leaf tissue inoculated with water [32] **(H')** *Zea mays* gene Zm00001eb015130, stalk mixture 3 days post inoculation with F. verticillioides wild type and fsr1 mutant [33] **(I')** *Zea mays* gene Zm00001eb110060, 63 day old root tissue treated with low nitrate for 24 hours [34] **(J')** *Zea mays* gene Zm00001eb132390, 15 days after planting endosperm tissue [35].

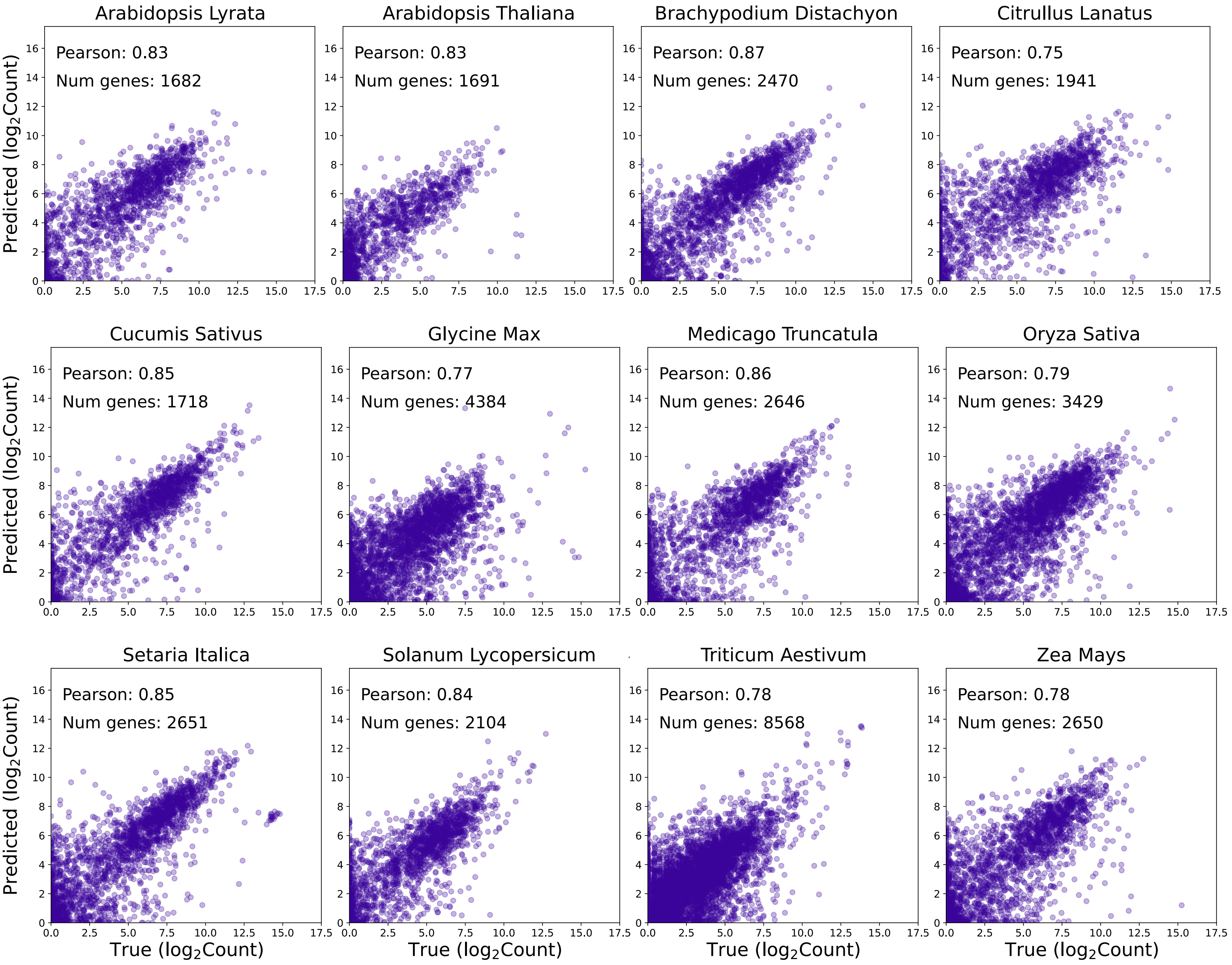


**Figure S2 - Examples of scatter plots of predicted vs true gene expression log2(count) values for validation set genes.** Each plot is for an individual species and an individual experimental condition. **(A)** *Arabidopsis lyrata* pistil tissue 3 days after being infected with fusarium graminearum [1] **(B)** *Arabidopsis thaliana* 5-6 week old male meiocyte [2] **(C)** *Brachypodium distachyon* tissue from an emerging 3rd leaf 13 days after germination, collected after grown in continuous light and continuous 28 °C temperature for 15.5 hours [5] **(D)** *Citrullus lanatus* young-ripening fruit 30 days after pollination [6] **(E)** *Cucumis sativus* 1 month old tendril tissue [8] **(F)** *Glycine max* embryo 45 days post initial seed (3-4mm) filling [11-15] **(G)** *Medicago truncatula* root 7 days after *Rhizoctonia solani* AG8 inoculation [16] **(H)** *Oryza sativa* leaf tissue at the 4 leaf stage [19] **(I)** *Setaria italica* L1 tiller bud 12 days after planting [23] **(J)** *Solanum lycopersicum* mature leaf tissue **(K)** *Triticum aestivum* seedling at the 3 leaf stage treated with ABA for 6 hours [30] **(L)** *Zea mays* stalk mixture 3 days post inoculation with *F. verticillioides* wild type and *fsr1* mutant [33].

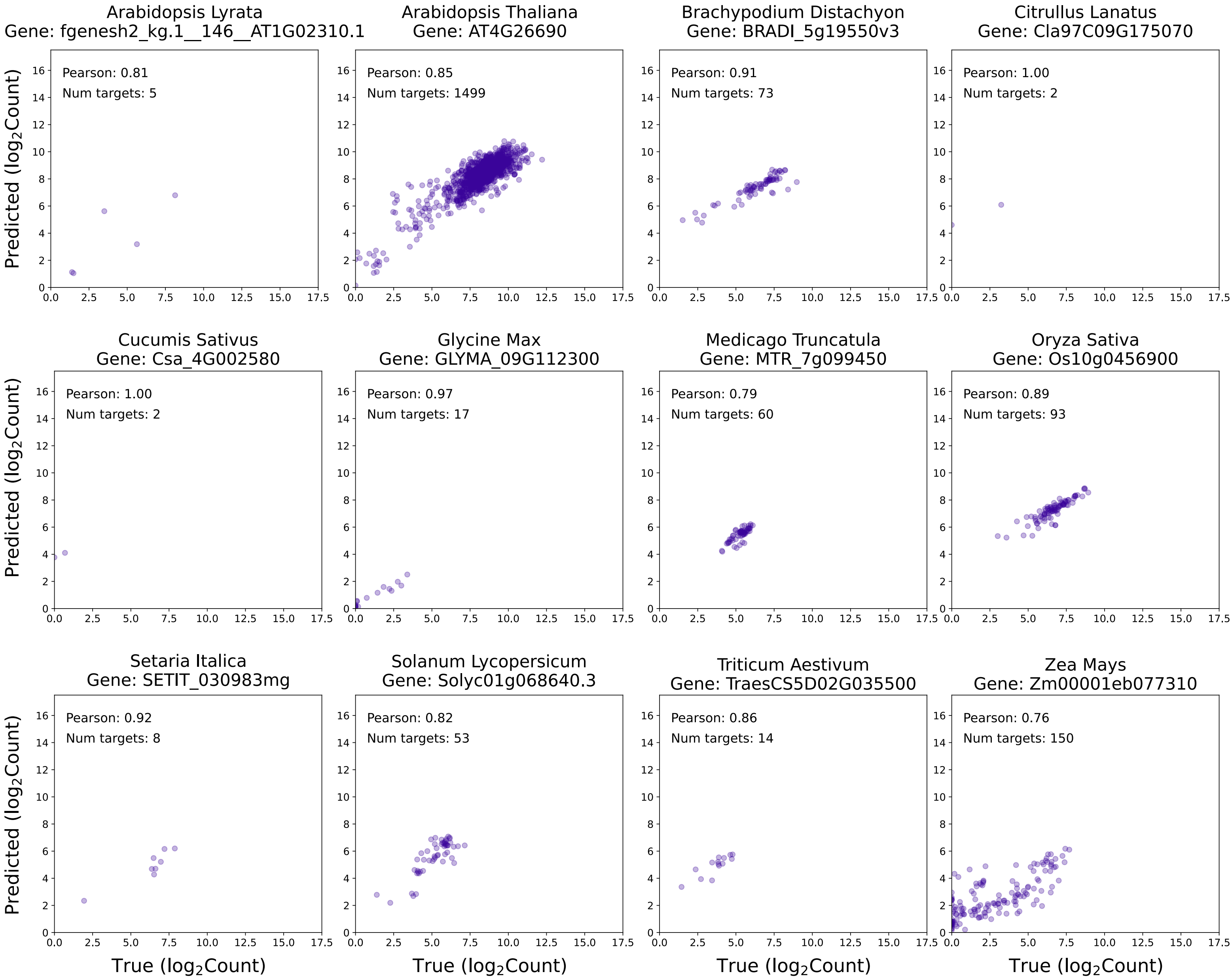


**Figure S3 - Examples of Scatter plots of predicted vs true gene expression**.
The $\log_2$(count) values for an individual gene in an individual species across all experimental RNA-seq conditions

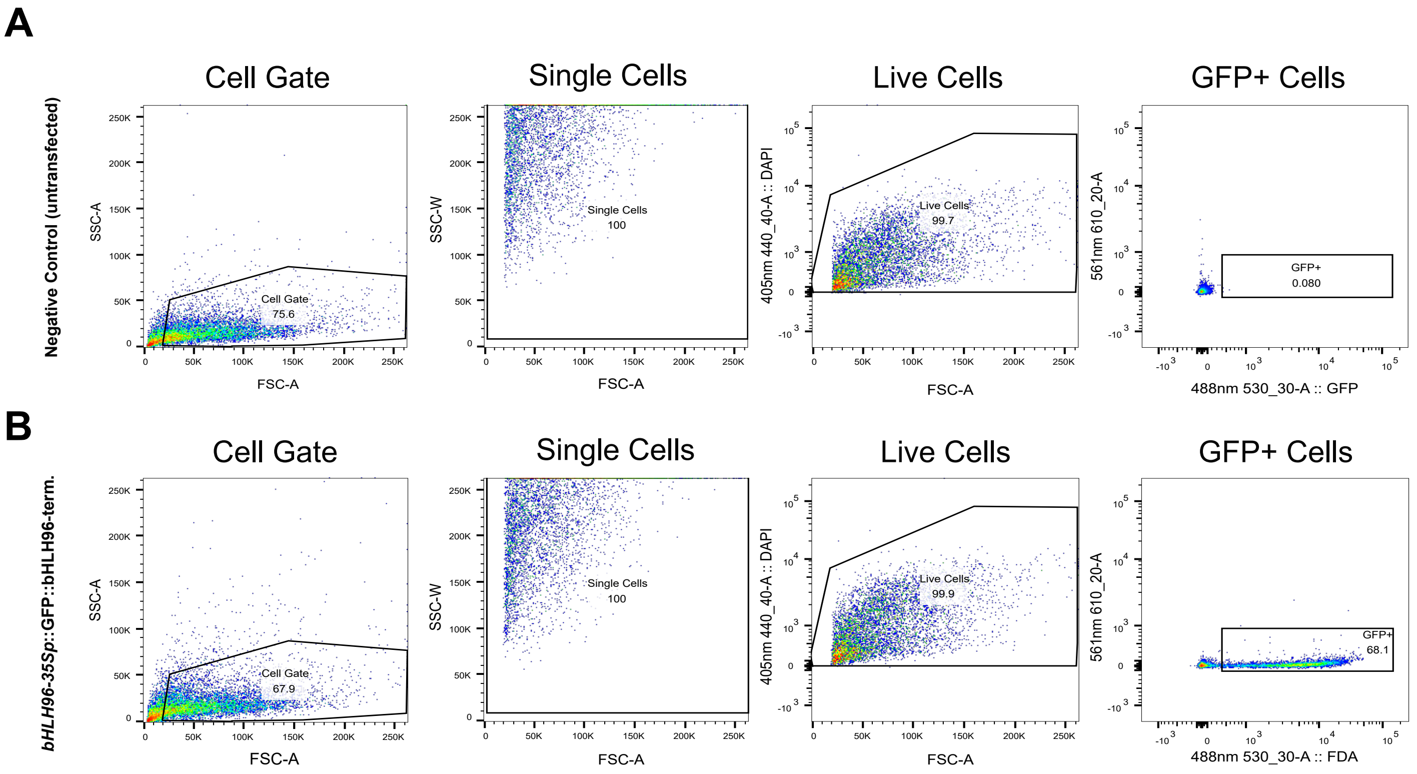


**Figure S4 - Gating boundaries for FACS-sorted cells (set with FlowJo).**
**(A)** Untransfected cells (negative control) and **(B)** positive control (*bHLH96p - 35S*::GFP) with gating in subsequent boundaries for: cell material, single cells (to remove doublets), living cells and define windows for GFP+ cells. FSC = forward scatter, SSC = side scatter, 488nm 530_30-A::FDA = GFP channel, 561nm 610_20-A mCherry channel (for comparison purposes).

#### Supplementary References


1. Accessed at: https://www.ncbi.nlm.nih.gov/bioproject/PRJNA408031

2. Warmerdam S, Sterken MG, Van Schaik C, Oortwijn MEP, Lozano-Torres JL, Bakker J, Goverse A, Smant G. Mediator of tolerance to abiotic stress ERF6 regulates susceptibility of Arabidopsis to Meloidogyne incognita. Mol Plant Pathol. 2019 Jan;20(1):137-152. doi: 10.1111/mpp.12745. Epub 2018 Oct 24. PMID: 30160354; PMCID: PMC6430479.

3. Trejo-Arellano MS, Mehdi S, de Jonge J, Dvorák Tomastíková E et al. Dark-Induced Senescence Causes Localized Changes in DNA Methylation. *Plant Physiol* 2020 Feb;182(2):949-961. PMID: 31792150

4. Szaker HM, Darkó É, Medzihradszky A, Janda T, Liu HC, Charng YY, Csorba T. miR824/AGAMOUS-LIKE16 Module Integrates Recurring Environmental Heat Stress Changes to Fine-Tune Poststress Development. Front Plant Sci. 2019 Nov 25;10:1454. doi: 10.3389/fpls.2019.01454. PMID: 31824525; PMCID: PMC6886564.

5. Accessed at: https://www.ncbi.nlm.nih.gov/bioproject/PRJEB32498

6. Accessed at: https://www.ncbi.nlm.nih.gov/bioproject/PRJNA564186

7. Accessed at: https://www.ncbi.nlm.nih.gov/bioproject/PRJNA563969

8. Accessed at: https://ncbi.nlm.nih.gov/bioproject/PRJNA531735

9. Zhang Z, Wang B, Wang S, Lin T, Yang L, Zhao Z, Zhang Z, Huang S, Yang X. Genome-wide Target Mapping Shows Histone Deacetylase Complex1 Regulates Cell Proliferation in Cucumber Fruit. Plant Physiol. 2020 Jan;182(1):167-184. doi: 10.1104/pp.19.00532. Epub 2019 Aug 4. PMID: 31378719; PMCID: PMC6945849.

10. Accessed at: https://www.ncbi.nlm.nih.gov/bioproject/PRJNA531735

11. Aghamirzaie D, Nabiyouni M, Fang Y, Klumas C et al. Changes in RNA Splicing in Developing Soybean (Glycine max) Embryos. *Biology (Basel)* 2013 Nov 21;2(4):1311-37. PMID: 24833227

12. Collakova E, Aghamirzaie D, Fang Y, Klumas C et al. Metabolic and Transcriptional Reprogramming in Developing Soybean (Glycine max) Embryos. *Metabolites* 2013 May 14;3(2):347-72. PMID: 24957996

13. Aghamirzaie D, Batra D, Heath LS, Schneider A et al. Transcriptome-wide functional characterization reveals novel relationships among differentially expressed transcripts in developing soybean embryos. *BMC Genomics* 2015 Nov 14;16:928. PMID: 26572793

14. Collakova, Eva; Aghamirzaie, Delasa; Fang, Yihui; Klumas, Curtis; Tabataba, Farzaneh; Kakumanu, Akshay; Myers, Elijah; Heath, Lenwood S.; Grene, Ruth. Metabolic and Transcriptional Reprogramming in Developing Soybean (Glycine max) Embryos. Metabolites 2013, 3(2), 347-372; doi: 10.3390/metabo3020347

15. Aghamirzaie, Delasa; Nabiyouni, Mahdi; Fang, Yihui; Klumas, Curtis; Heath, Lenwood S.; Grene, Ruth; Collakova, Eva. 2013. Changes in RNA Splicing in Developing Soybean (Glycine max) Embryos. Biology 2013, 2(4), 1311-1337; doi:10.3390/biology204131

16. Accessed at: https://www.ncbi.nlm.nih.gov/bioproject/PRJEB19425

17. Accessed at: https://www.ncbi.nlm.nih.gov/bioproject/PRJEB42202

18. Accessed at: https://www.ncbi.nlm.nih.gov/bioproject/PRJNA590945

19. Accessed at: https://www.ncbi.nlm.nih.gov/bioproject/PRJNA515816

20. Bessho-Uehara K, Masuda K, Wang DR, Angeles-Shim RB, Obara K, Nagai K, Murase R, Aoki SI, Furuta T, Miura K, Wu J, Yamagata Y, Yasui H, Kantar MB, Yoshimura A, Kamura T, McCouch SR, Ashikari M. *Regulator of Awn Elongation 3*, an E3 ubiquitin ligase, is responsible for loss of awns during African rice domestication. Proc Natl Acad Sci U S A. 2023 Jan 24;120(4):e2207105120. doi: 10.1073/pnas.2207105120. Epub 2023 Jan 17. PMID: 36649409; PMCID: PMC9942864.

21. Song XJ, Kuroha T, Ayano M, Furuta T et al. Rare allele of a previously unidentified histone H4 acetyltransferase enhances grain weight, yield, and plant biomass in rice. *Proc Natl Acad Sci U S A* 2015 Jan 6;112(1):76-81. PMID: 25535376

22. Accessed at: https://www.ncbi.nlm.nih.gov/bioproject/PRJNA557819

23. Accessed at: https://www.ncbi.nlm.nih.gov/bioproject/PRJNA622604

24. Liu X, Tang S, Jia G, Schnable JC, Su H, Tang C, Zhi H, Diao X. The C-terminal motif of SiAGO1b is required for the regulation of growth, development and stress responses in foxtail millet (Setaria italica (L.) P. Beauv). J Exp Bot. 2016 May;67(11):3237-49. doi: 10.1093/jxb/erw135. Epub 2016 Apr 4. PMID: 27045099; PMCID: PMC4892719.

25. Liu X, Tang S, Jia G, Schnable JC, Su H, Tang C, Zhi H, Diao X. The C-terminal motif of SiAGO1b is required for the regulation of growth, development and stress responses in foxtail millet (Setaria italica (L.) P. Beauv). J Exp Bot. 2016 May;67(11):3237-49. doi: 10.1093/jxb/erw135. Epub 2016 Apr 4. PMID: 27045099; PMCID: PMC4892719.

26. Accessed at: https://www.ncbi.nlm.nih.gov/bioproject/PRJNA339202

27. Liu L, Lu Y, Wei L, Yu H et al. Transcriptomics analyses reveal the molecular roadmap and long non-coding RNA landscape of sperm cell lineage development. *Plant J* 2018 Oct;96(2):421-437. PMID: 30047180

28. Xia C, Huang J, Lan H, Zhang C. Long-Distance Movement of Mineral Deficiency-Responsive mRNAs in *Nicotiana Benthamiana*/Tomato Heterografts. Plants (Basel). 2020 Jul 10;9(7):876. doi: 10.3390/plants9070876. PMID: 32664315; PMCID: PMC7412313.

29. Xia C, Zheng Y, Huang J, Zhou X, Li R, Zha M, Wang S, Huang Z, Lan H, Turgeon R, Fei Z, Zhang C. Elucidation of the Mechanisms of Long-Distance mRNA Movement in a *Nicotiana benthamiana*/Tomato Heterograft System. Plant Physiol. 2018 Jun;177(2):745-758. doi: 10.1104/pp.17.01836. Epub 2018 May 2. PMID: 29720554; PMCID: PMC6001325.

30. Accessed at: https://www.ncbi.nlm.nih.gov/bioproject/PRJNA626525

31. Accessed at: https://www.ncbi.nlm.nih.gov/bioproject/PRJEB21835

32. Accessed at: https://www.ncbi.nlm.nih.gov/bioproject/PRJNA485724

33. Accessed at: https://www.ncbi.nlm.nih.gov/bioproject/PRJNA423270

34. Accessed at: https://pubmed.ncbi.nlm.nih.gov/26198256/

35. Zhan J, Li G, Ryu CH, Ma C et al. Opaque-2 Regulates a Complex Gene Network Associated with Cell Differentiation and Storage Functions of Maize Endosperm. *Plant Cell* 2018 Oct;30(10):2425-2446. PMID: 30262552
